# Co-delivered PD-L1 rescues the protective efficacy mediated by an AAV-expressed HIV-1 bNAb

**DOI:** 10.64898/2026.05.29.728706

**Authors:** Michael Kuipa, Abubakarr A. Koroma, Isai Leguizamo, Priya Dhole, Yash Barot, Michelle Y.-H. Lee, Gregory K. Tharp, Shan Liang, Magdalen Chouinard, Stephanie Ehnert, Stacey Weissman, Casey Whitehead, Rachelle L. Stammen, Jennifer S. Wood, Elizabeth H. Curran, Deepa Machiah, Evan D. Dessasau, Yoshiaki Nishimura, Jun Xie, Guangping Gao, Sumit Verma, Deanna A. Kulpa, Ian N. Moore, Steven E. Bosinger, Matthew R. Gardner

## Abstract

Adeno-associated virus (AAV)-delivered anti-HIV-1 broadly neutralizing antibodies (bNAbs) could prevent and treat HIV-1 infection but are limited by host immune responses, specifically anti-drug antibodies (ADA). We tested whether PD-L1-mediated immune shielding could improve the consistency of AAV-delivered bNAb expression from muscle tissue in rhesus macaques. AAV9.PD-L1 co-delivery with AAV9.3BNC117 reduced the occurrence of ADA and T cell responses and improved the durability of 3BNC117 expression for one year post administration. Importantly, 5 of 6 macaques that received co-delivered AAV9.PD-L1 vectors were protected against ten repeated SHIV_AD8-EO_ challenges. Histopathological and spatial transcriptomic profiling showed that AAV9.PD-L1 co-delivery prevented severe local inflammation, muscle injury, and tertiary lymphoid structure formation at the administration site. Thus, immune shielding could serve as a strategy to prolong transgene expression from muscle-directed AAV-delivered biologics.

## Introduction

HIV-1 broadly neutralizing antibodies (bNAbs) offer an alternative to antiretroviral therapy (ART) for HIV-1 treatment and prevention (*1–5*) but require repeated in-clinic administration to sustain protective concentrations. Adeno-associated virus (AAV)-mediated delivery of bNAb genes to long-lived skeletal muscle cells could enable sustained antibody production and years-long protection from a single administration (*6*). However, bNAbs are inherently immunogenic due to their high levels of somatic hypermutation (SHM) (*7*). Since potency and breadth often correlate with the extent of SHM, the antibodies most desirable for AAV-based HIV-1 applications tend to be the most immunogenic. Consequently, bNAbs often trigger strong anti-drug antibody (ADA) responses against the bNAb variable regions, causing concentrations to decline within 2–8 weeks post administration (*7–13*).

The ADA barrier has been exemplified by our group and others through the delivery of 3BNC117. This CD4-binding site bNAb has SHM rates around 28% for the heavy chain and 20% for the light chain, much higher than the typical 6% rate seen in the memory B cell repertoire. Thus, it is not surprising that 3BNC117 usually has higher ADA responses compared to other bNAbs tested. Successful sustained expression of 3BNC117 is therefore a stringent benchmark for strategies to overcome host immune responses to immunogenic transgenes.

The PD-1/PD-L1 immune checkpoint pathway is a well-characterized mechanism by which PD-L1-expressing cells can suppress immune cell activity and promote immune tolerance through PD-1 signaling (*14*). While the PD-1/PD-L1 immune checkpoint pathway has been leveraged for cancer immunotherapy, we hypothesized that AAV-expressed PD-L1 could provide a similar evasion mechanism from the host immune response as seen in certain cancers. In this study we evaluated the use of PD-L1 as an immune shielding strategy to sustain transgene expression of the immunogenic bNAb, 3BNC117, in a nonhuman primate (NHP) model.

## Results

### Co-delivered PD-L1 significantly increases 3BNC117 concentrations while significantly reducing ADA in rhesus macaques

To determine whether co-delivered PD-L1 would overcome host immune responses against an immunogenic transgene, we first designed AAV9 vectors encoding 3BNC117 and rhesus PD-L1 (**Fig. S1**). Three groups of rhesus macaques (n=6 each) received AAV9.3BNC117 with or without AAV9.PD-L1, or AAV9.PD-L1 only as a control (hereafter denoted as 3BNC117-only, 3BNC117 plus PD-L1, or PD-L1-only) (**Fig. 1A**). AAV9 vectors were well tolerated, with no reported adverse events, and all macaques gained weight at the expected rate for the 52 weeks (**Fig. S2A-C**). ELISA quantification of serum 3BNC117 concentrations revealed two distinct expression profiles, which we termed “sustained” and “transient”. In the sustained profile, 3BNC117 concentrations plateaued by ∼week 10 and remained ≥50 µg mL⁻¹ thereafter; a concentration exceeding that required for protection in macaques (*15*) and approaching concentrations considered necessary for therapy (*4, 5*). In the transient profile, concentrations rose initially but declined sharply between weeks 6–8 to low or undetectable amounts for an extended period.

**Figure 1.**
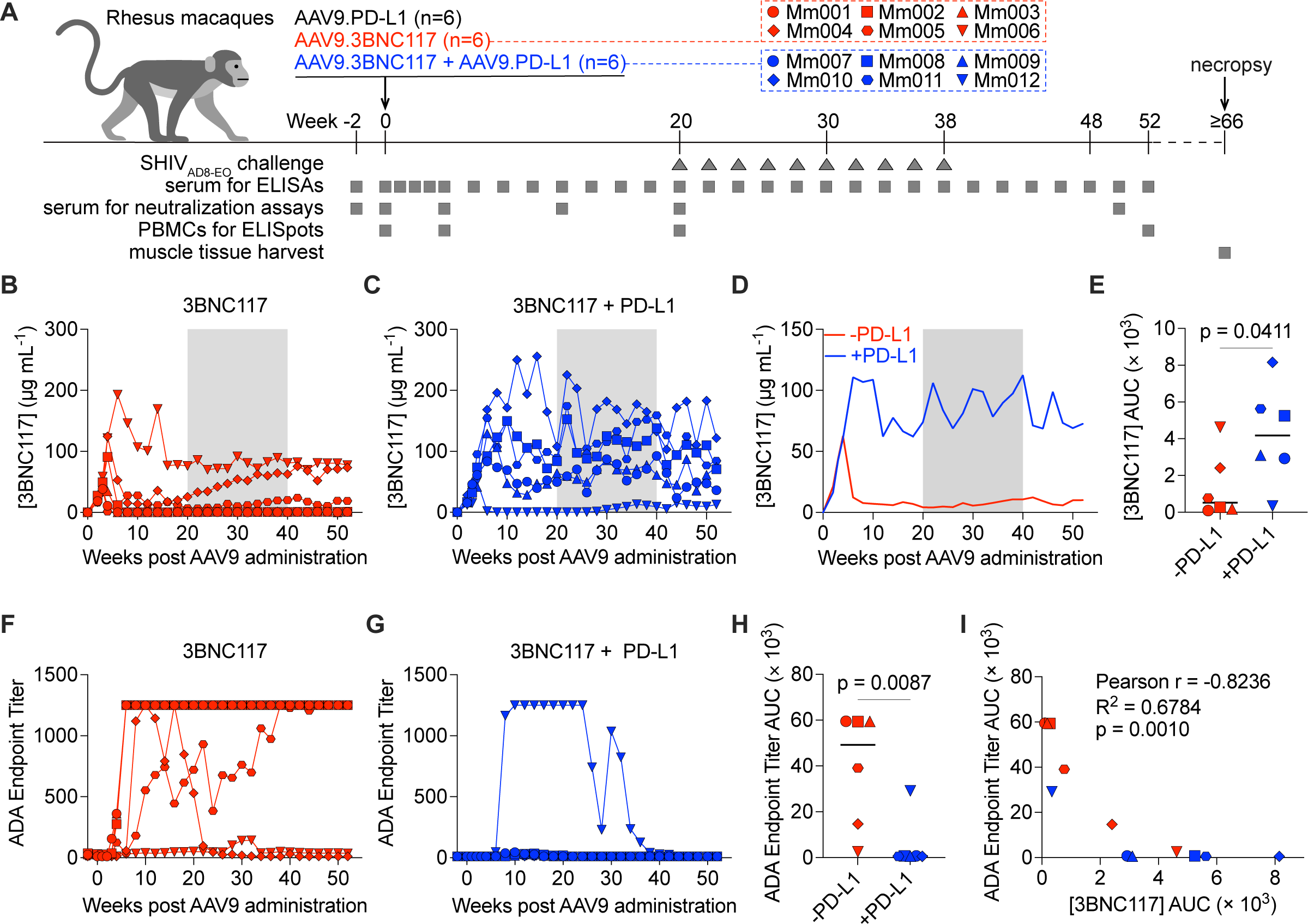
AAV-delivered bNAb expression and ADA in rhesus macaques when co-delivered with an AAV9.PD-L1 vector. **(A)** Overview of the study and sample collection. Three groups of rhesus macaques (n = 6 per group) received AAV vectors as indicated. Timeline indicates weeks post-AAV9 administration. SHIV_AD8-EO_ challenges were initiated at week 20. Samples were collected periodically and used for assays as indicated. Serum concentrations in individual macaques that received **(B)** AAV9.3BNC117 only or **(C)** AAV9.3BNC117 plus AAV9.PD-L1 as measured by gp120 ELISA or gp120 resurfaced stabilized core (RSC3) over 52 weeks. **(D)** Median serum 3BNC117 concentrations in macaques that received AAV9.3BNC117. **(E)** Comparison of average AUC for 3BNC117 serum concentrations. Serum 3BNC117 ADA endpoint titers in individual macaques that received **(F)** AAV9.3BNC117 only or **(G)** AAV9.3BNC117 plus AAV9.PD-L1 measured by anti-3BNC117 ELISA. **(H)** Comparison of average AUC for 3BNC117 ADA endpoint titers. **(I)** Pearson correlation plot of serum 3BNC117 concentration and 3BNC117 ADA endpoint titers in macaques that received AAV9.3BNC117. Pearson r value, R^2^ value, and p value (two-tailed t test), as determined by correlation analysis are included. Gray shading in panels (B–D) indicates the SHIV_AD8-EO_ challenge phase. Bars in (E) and (H) represent the median. Symbols in (B–I) correspond to those in (A). Statistical significance in (E) and (H) was determined by two-tailed Mann-Whitney test and defined as p ≤ 0.05.

In the 3BNC117-only group, all but one macaque (Mm006) demonstrated a transient expression profile, three of which (Mm001, Mm002 and Mm003) resulted in concentrations below the limit of detection for the remainder of the study (**Fig. 1B**). In contrast, 4 of 6 macaques in the 3BNC117 plus PD-L1 group had sustained expression profiles, with average concentrations ranging from 58–149 µg mL^−1^ (**Fig. 1C**). Of the remaining two macaques, Mm007 had a near-sustained profile, with concentrations reaching ≥50 µg mL⁻¹ mid-study (average 54 µg mL^−1^) and Mm012 had a transient profile, averaging 7 µg mL^−1^. Overall, median concentrations were ∼18-fold higher with PD-L1 co-delivery (**Fig. 1D**) and AUC differed significantly between the two groups (p= 0.0411) (**Fig. 1E**). Collectively, these data demonstrate that co-delivered PD-L1 rescues expression of AAV-delivered 3BNC117, resulting in measurable serum concentrations for over a year.

### PD-L1 co-delivery prevents ADA responses against AAV-expressed bNAbs

Due to the success with durable 3BNC117 expression when co-delivered with PD-L1, we next assessed whether these results were from decreased host immune responses. To assess if PD-L1 reduces ADA responses, we measured ELISA reactivity against purified 3BNC117 (**Fig. 1F,G**) in serum over 52 weeks. In the 3BNC117-only group, 5 of 6 macaques developed ADA responses. The three macaques with clear transient 3BNC117 expression profiles (Mm001, Mm002, Mm003) developed high, sustained ADA responses (**Fig. 1F, Fig. S3A**). The two macaques (Mm004 and Mm005) with transient 3BNC117 expression that rebounded later in the study, showed a transient ADA response in Mm004 and delayed ADA response in Mm005. Mm006, the macaque with sustained 3BNC117 expression in the group, had minimal ADA titers above baseline (**Fig. 1F, Fig. S3A**). ADA responses were detectable by week 8, coinciding with loss of bNAb expression and there was a trend of higher ADA endpoint titers in macaques with lower bNAb concentrations (**Fig S3A-D**).

In the 3BNC117 plus PD-L1 group, we observed only 1 of 6 macaques with an ADA response (**Fig. 1G, Fig. S3B**). Mm012 developed a clear ADA response, aligning with its transient bNAb expression profile (**Fig. 1G, Fig. S3B**). ADA endpoint titer AUCs were significantly lower in the 3BNC117 plus PD-L1 group compared with the 3BNC117-only group (p = 0.0087) (**Fig. 1F**). We also observed a significant negative correlation between 3BNC117 concentration AUCs and ADA endpoint titers (p = 0.0010) (**Fig. 1I**). However, when assessing anti-AAV9 capsid response, all 18 macaques developed a neutralizing anti-AAV9 antibody response (**Fig. S4A-D**). There were no differences in anti-AAV9 antibody titer AUCs between groups (**Fig. S4D**). Together, these data show that co-delivered PD-L1 may only affect the antibody response against the expressed transgene and not against the AAV capsid.

### PD-L1 co-delivery attenuates CD8+ T cell mediated immunity against 3BNC117

We then assessed anti-bNAb cellular immunity using IFNγ ELISpot assays on week-4 and week-52 PBMCs (**Fig. S4E,F**). PBMC reactivity was most consistently directed against the 3BNC117 VarH peptide pool with a subset responding against the 3BNC117 VarL peptide pool (**Fig. S4E**). We noted significant anti-3BNC117 VarH peptide pool reactivity in the 3BNC117 only group compared to the 3BNC117 plus PD-L1 group at week 52 (p = 0.0022) (**Fig. S4F**). Macaques Mm002 and Mm003 in the 3BNC117-only group developed strong reactivity to multiple peptide pools. Thus, co-delivered PD-L1 additionally reduces T cell activation against expressed 3BNC117 peptides underscoring its beneficial impact on the expressed transgene.

### AAV-expressed bNAbs remain functional *ex vivo* and exhibit neutralization breadth

We next assessed the serum neutralization activity of the AAV-delivered 3BNC117 in all macaques at 20 weeks post administration. Sera were tested against 11 HIV-1 pseudoviruses, SHIV_AD8-EO_ pseudovirus (Env of challenge virus), and SIVmac239 (negative control) using a TZM-bl neutralization assay. As expected, macaques with measurable 3BNC117 (**Fig. 2A,B**) serum concentrations neutralized pseudoviruses sensitive to the bNAb. Median ID₅₀ (infectious dose 50%) titers were higher in the PD-L1-treated group, consistent with higher bNAb concentrations determined by ELISA. Serum from macaques in the PD-L1-only group exhibited no neutralizing activity (**Fig. 2C**). As anticipated, serum bNAb concentrations correlated strongly with SHIV_AD8-EO_ ID₅₀ neutralization titers across groups (p < 0.0001) (**Fig. 2D**). These data support the feasibility of achieving broad, durable HIV-1 protection through a single vectored administration, with neutralization breadth and potency that have remained elusive for HIV-1 vaccines.

**Figure 2.**
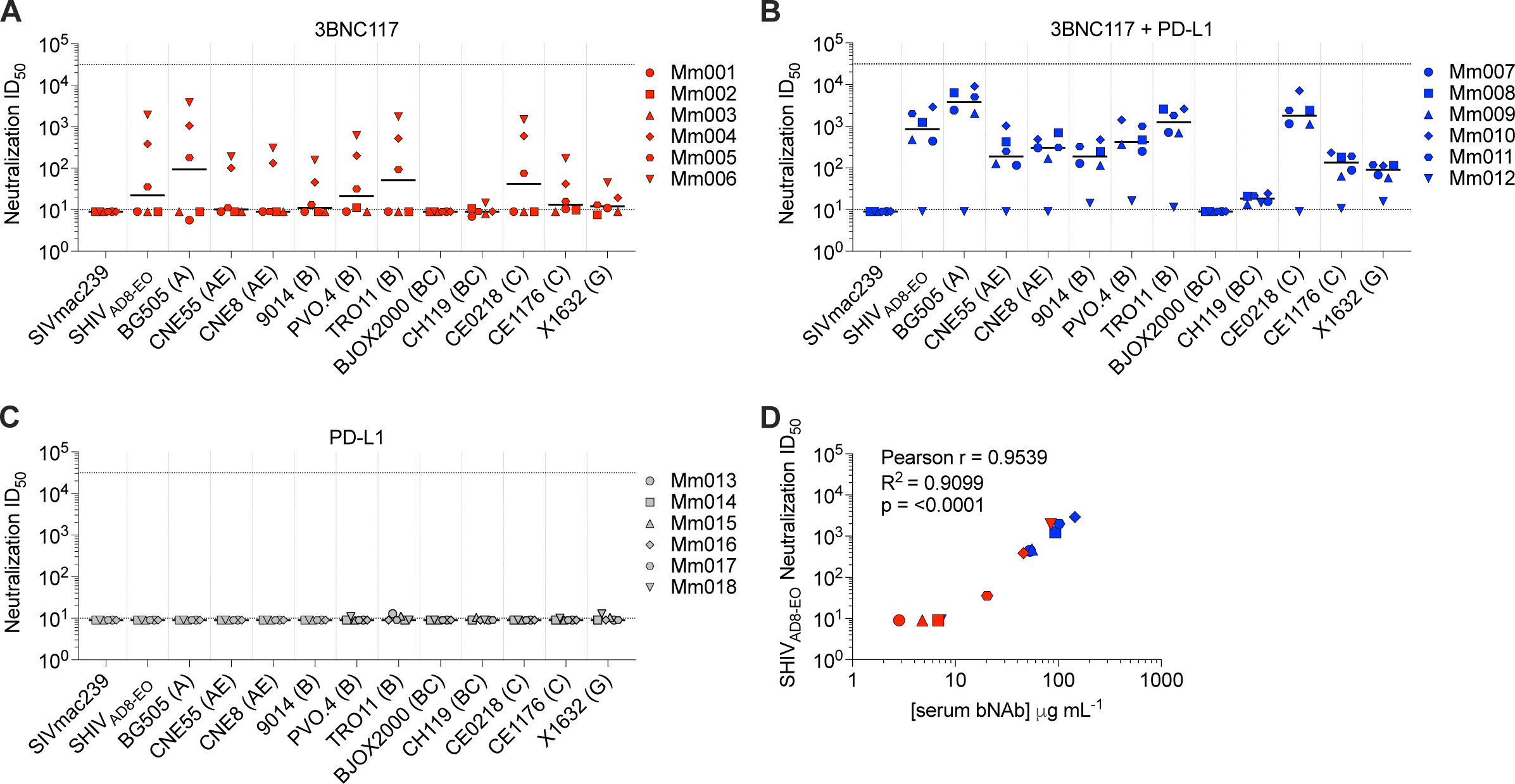
*Ex vivo* neutralization breadth of AAV-expressed 3BNC117. Week-20 serum from all 18 macaques was tested for neutralization against a panel of 11 HIV-1 pseudoviruses, the challenge virus SHIV_AD8-EO_ pseudovirus, and SIVmac239, using a TZM-bl neutralization assay. ID₅₀ values for each macaque per group are reported for the indicated pseudovirus and those samples that did not reach 50% neutralization were normalized to a value of <10. Week-20 serum ID₅₀ titers in individual macaques that received **(A)** AAV9.3BNC117 only, **(B)** AAV9.3BNC117 plus AAV9.PD-L1 or **(C)** AAV9.PD-L1 only. In (A–C) HIV-1 clade is indicated in parentheses, and black lines denote median ID₅₀ values. **(D)** Pearson correlation plot of serum bNAb concentration as determined in Fig. 1 and SHIV_AD8-EO_ ID₅₀ neutralization titer in macaques that received AAV9.3BNC117. Note symbols in (D) match those used in panels (A–C). Pearson r value, R^2^ value, and p value (two-tailed t test), as determined by correlation analysis are included. Statistical significance is defined as p value ≤ 0.05.

### Co-delivery of PD-L1 rescues the protective efficacy mediated by AAV-expressed 3BNC117

To evaluate the protective efficacy of the AAV-delivered bNAbs, we conducted a low-dose intrarectal SHIV challenge study. In the PD-L1-only control group, the median number of challenges to infection was 4.5 (**Fig 3A,B**), with peak plasma viral loads (PVLs) ranging from 8.03 × 10⁵ to 3.93 × 10⁷ copies mL^−1^ (**Fig. 3B**). In the 3BNC117-only group, 4 of 6 macaques were infected after one challenge, with serum 3BNC117 concentrations ranging from undetectable to ∼7 µg mL^−1^ at the time of infection (**Fig. 3A**). Peak PVLs in the infected macaques ranged from 2.08 to 6.57 × 10⁶ copies mL^−1^ (**Fig. 3C**). The remaining two macaques (Mm004, Mm006) resisted all 10 challenges. In contrast, 5 out 6 macaques in the 3BNC117 plus PD-L1 group resisted all 10 challenges which was significantly different compared to the PD-L1-only group (p = 0.0078) (**Fig. 3A**). Additionally, we observed a trend towards significance in improved protection compared to the 3BNC117-only group (p = 0.0673). Mm012 in the 3BNC117 plus PD-L1 group was infected after two challenges, with undetectable serum 3BNC117 concentrations at the time of infection. Peak PVL reached 5.37 × 10⁷ copies mL^−1^ (**Fig. 3D**). These data highlight the utility of PD-L1 co-delivery for an AAV-expressed transgene to achieve its desired therapeutic outcome after a single vector administration.

**Figure 3.**
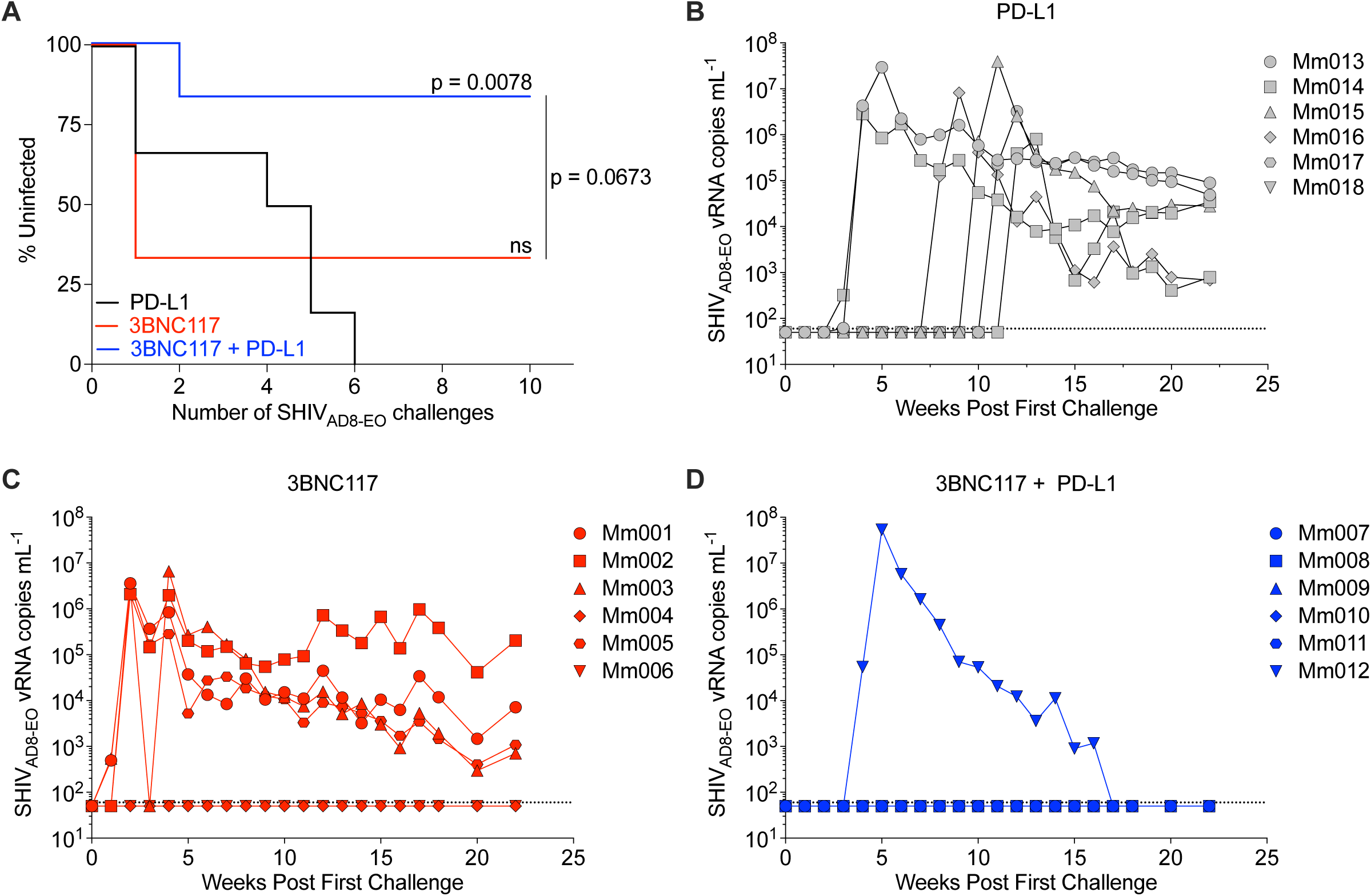
AAV-delivered bNAbs mediate protection from low-dose, intrarectal SHIV_AD8-EO_ challenges. **(A)** Kaplan-Meier analysis of protection from SHIV_AD8-EO_ infection over the course of ten biweekly intrarectal challenges (10 TCID₅₀ per exposure) in animals that received AAV9.3BNC117 compared to the control group, which received AAV9.PD-L1 only. Plasma SHIV_AD8-EO_ viral RNA mL^−1^ levels during the SHIV_AD8-EO_ challenge phase over time in macaques that received **(B)** AAV9.PD-L1 only, **(C)** AAV9.3BNC117 only, or **(D)** AAV9.3BNC117 plus AAV9.PD-L1. Viral loads were measured by qRT-PCR with a limit of detection of 60 copies mL^−1^ (dotted line). Statistical significance was determined by the Mantel-Cox test. ns, not significant.

### PD-L1 co-delivery prevents the development of tertiary lymphoid structures at the site of administration

Having observed correlations of 3BNC117 expression and ADA induction, we asked whether the local environment at the administration site might explain these results. We therefore analyzed hematoxylin and eosin (H&E)-stained quadriceps muscle sections collected near the site of AAV administration from all 18 macaques (**Fig. S5A-C**). We developed a five-point inflammation severity scoring system ranging from none/not overt (−) to marked (+++), representative images of which are shown in **Fig. 4A**. Notably, in the PD-L1-only group, 4 of 6 macaques had a “+/−” or “+” score, indicating that AAV9-delivered PD-L1 alone could induce low-grade inflammation. Additionally, in the 3BNC117 plus PD-L1 group, inflammation did not exceed a “+” grade. In contrast, inflammation was more severe in the 3BNC117-only group, with 4 of 6 macaques graded “++” or higher. Of note, two 3BNC117-only group macaques (Mm001 and Mm005) had marked inflammation (+++) (**Fig. 4B**). On examining how ADA responses related to local tissue inflammation, we observed a trend in macaques with greater inflammation exhibiting higher ADA responses, indicating that ADA may be a surrogate for the health of the local tissue environment.

**Figure 4.**
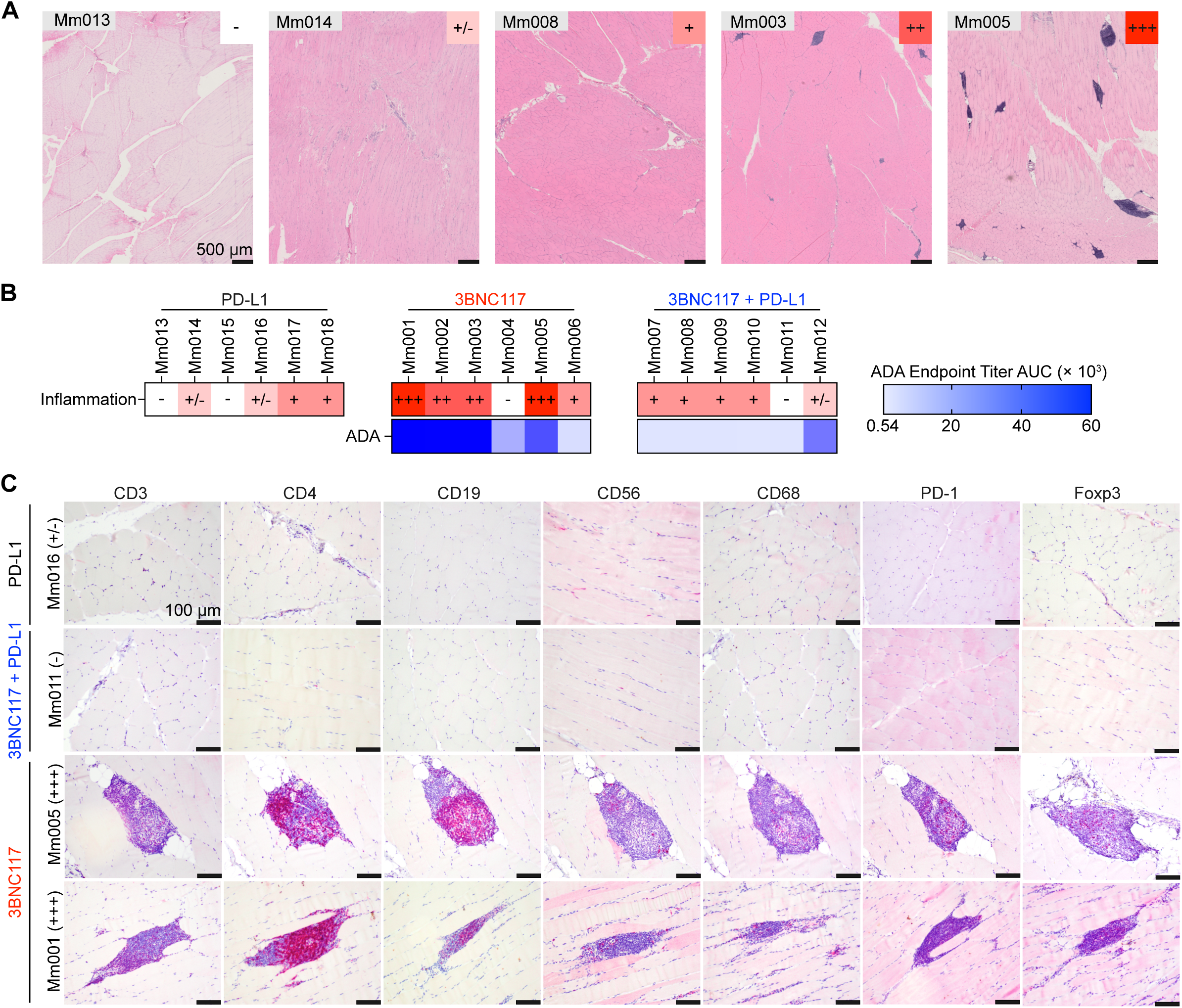
Inflammation assessment of muscle tissue at the site of AAV9 administration. Muscle tissue from the upper left quadriceps of all 30 macaques was harvested at necropsy. H&E stained FFPE muscle tissue sections were scored for inflammation. **(A)** Representative images demonstrating the tiered scoring system of inflammation. Scale bars = 500 µm. **(B)** Muscle inflammation severity scoring for all 18 macaques compared to ADA endpoint titer AUC. Darker colors indicate higher inflammation (red) or ADA endpoint titer (blue). **(C)** IHC staining for CD3, CD4, CD19, CD56, CD68, PD-1, and FoxP3 in representative animals with less severe inflammation (Mm011 and Mm016) versus those with more severe inflammation (Mm001 and Mm005). Scale bars = 100 µm. Inflammation scoring legend: -, no evidence of interstitial hypercellularity or overt inflammation; +/−, slight interstitial hypercellularity or minimal focal inflammatory cell infiltration; +, moderate interstitial hypercellularity or mild, diffuse inflammation; ++, evident inflammation with small-to-moderate foci showing multifocal distribution; +++, marked inflammation with moderate-to-large foci displaying multifocal and coalescing distribution. Severity scores were adjusted to accommodate changes to normal muscle fiber morphology such as muscle degeneration and/or necrosis.

Extensive inflammatory cell foci observed in macaques with severe inflammation (++ and +++) prompted immunohistochemistry (IHC) analysis of a subset of tissue samples for immune cell markers CD3, CD4, CD19, CD56, CD68, PD-1 and Foxp3 (**Fig. 4C**). Mm016 (PD-L1-only) and Mm011 (3BNC117 plus PD-L1) were selected for comparison. We identified T cells, B cells, NK cells, and macrophages in the samples with severe inflammation (Mm001 and Mm005) but not in those without (Mm011 and Mm016). These regions with inflammatory cell foci were consistent with the microanatomy of tertiary lymphoid structures (TLSs), ectopic lymphoid organs observed in pathological contexts (*16*). Of interest, for Mm005, we observed a germinal center-like structure of concentrated, yet segregated, CD4 and CD19 markers, a hallmark of a mature TLS. We also observed PD-1 and FoxP3 expression within the foci-like structures, potentially highlighting the presence of T follicular regulatory (Tfr) cells, T follicular helper (Tfh) cells, and/or T regulatory (Treg) cells. Additional examples of the TLSs are shown in **Fig. S6A,B** for Mm001 and Mm005. Based on H&E staining, all four macaques (Mm001, Mm002, Mm003, Mm005) with “++” or “+++” inflammation scores had structures resembling TLSs (**Fig. S5B,C**). All macaques with sustained ADA responses were found to have TLSs present in the muscle tissue. Importantly, we did not observe TLSs in the 12 macaques that received PD-L1 vectors. These data demonstrate the protective effects of PD-L1 co-delivery in reducing inflammation and TLS formation at the AAV administration site. Additionally, the development and persistence of TLSs in the muscle tissue suggests a mechanism for the sustained ADA response even after the bNAb is no longer detectable in the blood.

### Gene signatures confirm the TLSs found in the muscle tissue are actively producing antibodies and recruiting immune cells

To examine the immunopathological findings in greater detail, we performed Visium HD spatial transcriptomics on quadriceps muscle tissue biopsies from representative macaques in the 3BNC117-only group (Mm001) and the 3BNC117 plus PD-L1 group (Mm011). The 3BNC117-only biopsy featured multiple nuclei-dense pockets that were absent in the 3BNC117 plus PD-L1 biopsy (**Fig. 5A**). We found that B cell marker *MS4A1/CD20*, and T cell receptor (TCR) alpha and beta constant region gene segment (*TRAC*, *TRBC1*, *TRBC2*) transcripts were localized within discrete regions of the nuclei-dense pockets (**Fig. 5B**). *TRAC*, *TRBC2* transcript levels were >2000 barcodes, compared to <150 barcodes in the 3BNC117 plus PD-L1 biopsy (**Fig. 5C**). *MS4A1* and *TRBC1* transcript levels were >300 barcodes in the 3BNC117-only biopsy, while <10 barcodes for the 3BNC117 plus PD-L1 biopsy. Within the pocket-like structures of the 3BNC117-only biopsy, we observed localization of cyto/chemokine gene transcripts *CCL19*, *CCL21*, *LTA* and *LTB*, which are involved in TLS development (*17, 18*) (**Fig. 5D,E**). In contrast, TLS-related cyto/chemokine expression was minimal in the 3BNC117 plus PD-L1 biopsy. Likewise, *IGHM, JCHAIN* and *IGKC* transcripts (>8,500 barcodes each) were localized within the TLS-like pockets but were virtually absent in the 3BNC117 plus PD-L1 biopsy (<100 barcodes each) (**Fig. 5F,G**). Next, we assembled a TLS B cell signature comprising of *BANK1*, *CD22*, *BLK*, *SYK*, *CD79A*, and *CD79B* and found these transcripts (>3,000 barcodes total) to be localized within discrete regions of the immune pockets (**Fig. 5H**). Conversely, TLS B cell signature expression was found at low levels (<75 barcodes total) in the 3BNC117 plus PD-L1 biopsy (**Fig. 5I**).

**Figure 5.**
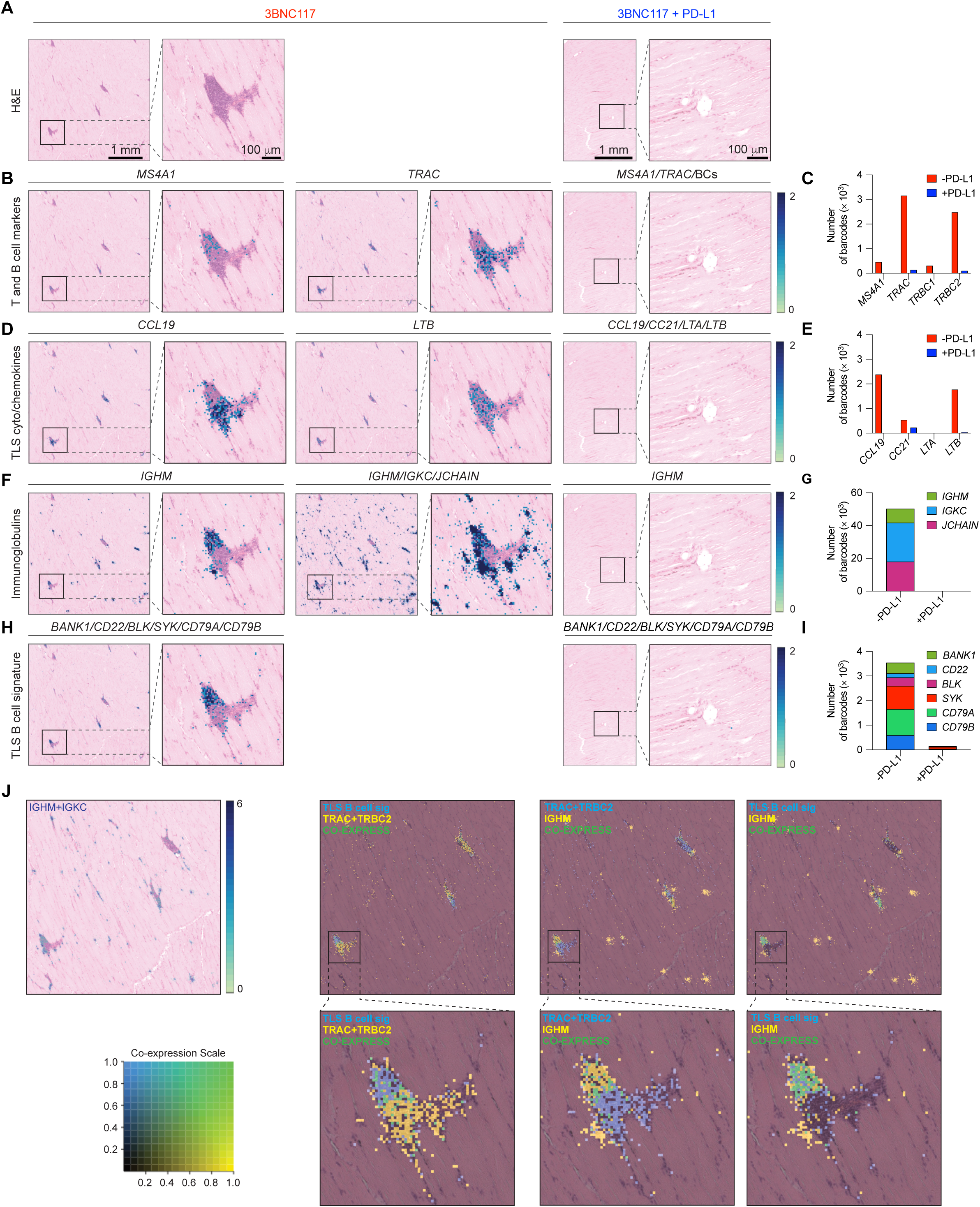
Transduction of muscle with co-delivered AAV9.PD-L1 reduces the formation of TLSs. **(A)** H&E staining of FFPE sections of rhesus macaque biopsies obtained from the upper left quadriceps muscle at necropsy (n = 1 for each condition) used for Visium HD spatial transcriptomics. Scale bars = 1 mm. The insets depict ROIs enlarged in the neighboring panel. Scale bars = 100 µm. **(B)** Localization of transcript expression of B cell marker *MS4A1* (CD20), and indicated TCR genes as single genes (3BNC117-only) or as a combination (3BNC117 plus PD-L1). **(D)** Spatially localized expression of indicated chemokines/cytokines regulating TLS formation as single genes (3BNC117-only) or as a combination (3BNC117 plus PD-L1). **(F)** Spatially localized expression of indicated immunoglobulin and J chain transcripts as individual genes or in combination. (**H)** Spatially localized expression of indicated TLS-signature genes in combination for each group. Transcript expression levels are depicted according to the log2 scale shown at right. **(C, E, G, I)** Sum of barcodes for individual transcripts over the entire spatial substrate. **(J)** Co-expression plots of indicated TCRs and TLS B cell genes with *IGHM*. The panel at top left depicts the expression of *IGHM* and *IGKC*. The co-expression color scale for each marker is depicted at bottom left. Individual marker colors are indicated on the inset of each panel.

To examine the anatomical organization of B and T lymphocytes within the pocket-like structures, we performed co-expression analyses of transcripts for *IGHM*, TCR and TLS B cell signature genes (**Fig. 5J**). The TLS-B cell signature had minimal co-localization with TCR (*TRAC*+*TRBC2*) transcripts. Similarly, *IGHM* and TCR expression was largely exclusive. In contrast, we observed a high degree of co-expression between TLS-B cell signature genes and *IGHM* (**Fig. 5J**). Collectively, these spatial transcriptomic analyses validate the formation and persistence of active TLSs at the AAV administration site in the absence of PD-L1 co-delivery.

### Co-delivery of PD-L1 protects muscle tissue from AAV-mediated injury

We next assessed spatial expression of genes associated with healthy muscle tissue, muscle regeneration, or active scarring/fibrosis defined recently in a Duchenne muscular dystrophy mouse model (*19*). In the 3BNC117-only biopsy, k-means clustering of genes produced clusters organized into two distinct zones defined by clusters 1 and 2, with a third cluster interspersed throughout (**Fig. 6A**). In the 3BNC117 plus PD-L1 biopsy, clusters had a high degree of spatial organization (**Fig. 6B**) and genes with the highest expression in clusters 1–3 were indicative of myocytes, with multiple myosins (*MYH1*, *MYL1*, *MYL2*, *MYBPC2*, *MYBPC1*, *MYOZ1*, *MYOT*), actin (*ACTA1*, *ACTN2*, *ACTN3*), titin (*TTN*), and troponins (*TNNC2*, *TNNT3*, *TNN12*) present in highly organized bundles (**Fig. 6D**). Although the top two clusters from the 3BNC117-only biopsy also had high myocyte-related gene expression (**Fig. 6D**), their spatial organization differed substantially from the 3BNC117 plus PD-L1 biopsy, with myosin-high clusters being less dense and more diffuse in their expression **(Fig 6A,B)**. Notably, we observed a cluster (C3) with high expression of immunoglobulin (*IGKC* and *IGHM*) and immune-related genes (*CD74*, *A2M*, *CD44*) that was absent in the 3BNC117 plus PD-L1 biopsy (**Fig. 6A,C**). We found that the 3BNC117 plus PD-L1 biopsy contained high expression of *MYH1* throughout, and expression of *MYL2* was organized into discrete bundles (**Fig. 6E-G**). In contrast, the 3BNC117-only biopsy contained a zone that was devoid of *MYH1* expression and contained disorganized patches of *MYL2* expression (**Fig. 6E,F**). Additionally, genes previously identified in muscle regeneration (*19*) (*MYH3*, *SPARC*, *IGFBP7* and *MYOG*) were expressed at much higher levels (>80,000 barcodes total) in the 3BNC117-only biopsy and were more highly enriched in the *MYH*1-free zone (**Fig. 6H-J**). In contrast, the 3BNC117 plus PD-L1 biopsy had <20,000 total barcodes of the regeneration signature genes and were predominantly *SPARC* and *MYOG* genes. Strikingly, expression of *MYH3*, a myosin found in embryonic and healing muscle, was highly enriched in the *MYH1*-free zone **(Fig. 6I)**. *MYH3* was expressed at high levels in the 3BC117-only biopsy (14,117 barcodes) but was virtually absent in the 3BNC117 plus PD-L1 biopsy (9 barcodes) (**Fig. 6J**). Lastly, genes associated with active fibrosis (*COL1A1*, *COL1A2*, *VIM*, *FN1*, *THBS4*) (*19*) were highly expressed in the *MYH1*-free regions of the 3BNC117-only biopsy, but not in the 3BNC117 plus PD-L1 biopsy (**Fig. 6K-M**). COL1A1, COL1A2, and VIM transcripts levels were >50,000 barcodes each in the 3BNC117-only biopsy while <16,000 barcodes each in the 3BNC117 plus PD-L1 biopsy. Additionally, FN1 and THBS4 transcripts levels were >9,000 barcodes each in the 3BNC117-only biopsy but <2,000 barcodes each in the 3BNC117 plus PD-L1 biopsy. Together, these data indicate that AAV-delivered 3BNC117 alone induces tissue injury that can be prevented by PD-L1 co-delivery.

**Figure 6.**
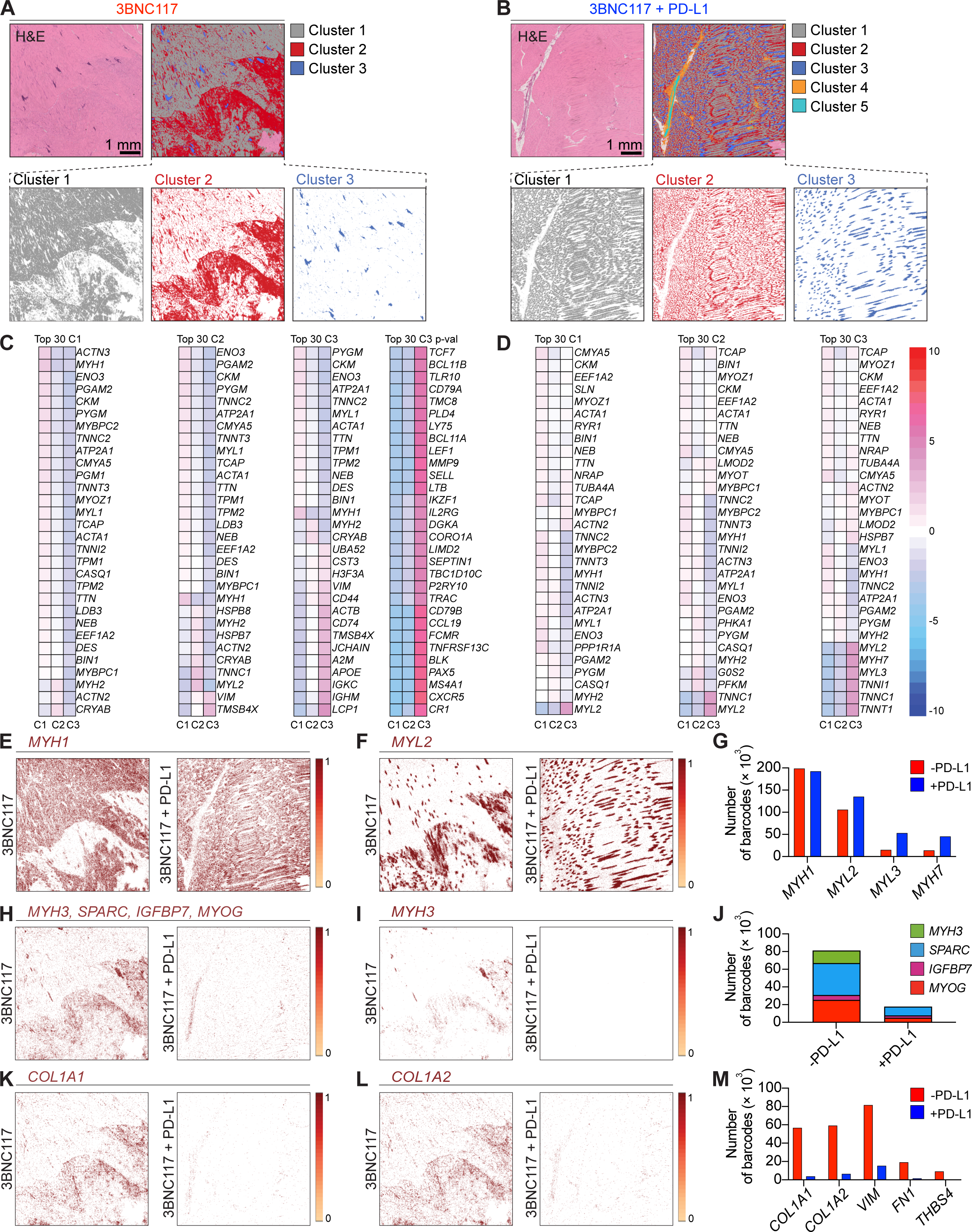
PD-L1 co-delivery protects muscle tissue from injury. H&E staining and k-means (k = 7) clustering of Visium HD spatial transcriptomics data from the upper left quadriceps biopsies from a representative macaque that received **(A)** AAV9.3BNC117 only or **(B)** AAV9.3BNC117 plus AAV9.PD-L1. Clusters containing <10 barcodes were omitted. The bottom panels show the three clusters with the highest barcode counts, each displayed individually in its assigned color. Heatmaps showing the highest 30 genes by average expression in each of the top three clusters from a representative macaque that received **(C)** AAV9.3BNC117 only or **(D)** AAV9.3BNC117 plus AAV9.PD-L1. The heatmap on the right in (C) shows the top 30 genes differentially expressed in Cluster 3 (C3) by p-value. Cluster numbers (C1-C3) are indicated at the bottom of each heatmap. The color scale depicted on the right indicates fold-change per gene of each cluster relative to other analyzed clusters on a log2 scale. **(E–G)** Spatial distribution and quantitation of indicated myosin genes found in healthy tissue. **(H–J)** Spatial distribution and quantitation of genes indicative of muscle regeneration. **(K–M)** Spatial distribution and quantitation of genes indicative of fibrosis. The color scale depicted at the right of spatial plots indicate barcode counts on a log2 scale.

## Discussion

Our findings establish that PD-L1-mediated immune shielding of transduced muscle enables durable expression of AAV-delivered 3BNC117 in rhesus macaques. A major barrier to evaluating expression and efficacy of AAV vectors in NHP preclinical models has been the host immune response against the transgene. We have overcome this hurdle through the co-delivery of PD-L1. As anticipated, we observed significantly higher 3BNC117 concentrations through lower host immune responses when we co-delivered an AAV vector expressing PD-L1. Our results further substantiate findings previously described only rodent models. Not only do these results provide a strategy for future evaluation of AAV vectors in NHPs, but they suggest translational applicability for improving AAV gene therapy in humans.

The limited durability of AAV-delivered bNAbs in preclinical NHP models and humans has been strongly associated with ADA responses, especially after intramuscular administration (*7–11*). Our results suggest TLS formation at the vector administration site as a previously unrecognized component of this process. TLSs are privileged immune niches for local antibody responses that typically form in pathological settings, including autoimmunity, viral infection, and tumors (*16*). In our model, they likely arise from chronic inflammation driven by low-level bNAb expression and/or the presence of AAV capsid at the administration site. While our initial hypothesis for using PD-L1 was to decrease immune cell effector functions on transduced muscle cells, our data demonstrate an unanticipated, important role of PD-L1 in preventing TLS formation at these administration sites. Notably, 4 of the 5 macaques in the 3BNC117-only group with high anti-3BNC117 ADA endpoint titers developed TLSs. In contrast, PD-L1 co-delivery resulted in 0 of 12 macaques developing TLSs. Our current findings add to prior work demonstrating the capacity of immune checkpoint molecule-based shielding to prevent deleterious immune responses, specifically the reduction of ADA, to sustain transgene expression (*20–22*).

Of clinical relevance, AAV-delivered bNAbs can provide durable virologic control, provided therapeutic concentrations are maintained (*6, 23*). In a rare, proof-of concept example, a SHIV_AD8-EO_-infected macaque that maintained concentrations of AAV1-delivered 10-1074 at >100 µg mL^−1^ and 3BNC117 at >50 µg mL^−1^ for more than three years, achieved complete virologic suppression throughout the study (*11*). In the present study, PD-L1 co-delivery enabled 5 of 6 macaques to sustain comparable 3BNC117 concentrations >50 µg mL^−1^ for a year post administration. While PD-L1 was associated with reduction in TLS formation, these findings underscore our previous work for improved antibody transgene expression through AAV capsid selection, cassette optimization, and vector administration strategies. Taken together, these findings suggest clinical translation for HIV therapy. 3BNC117 was used in this study for its high degree of somatic hypermutation and previously described immunogenicity. Thus, the PD-L1 strategy could offer similar results for any bNAb with lower degrees of somatic hypermutation.

Previous efforts to overcome immune responses against AAV-delivered transgenes have relied on systemic immunosuppression with calcineurin inhibitors (*9*), mTOR inhibitors (*24*), and corticosteroids (*25*), which require continued dosing and carry risks of toxicity and increased infection susceptibility. Other strategies for inducing transgene tolerance have been to target the liver (*26, 27*) and, recently, leveraging the tolerogenic neonatal immune system by administering AAV-delivered bNAbs at birth (*28*). We demonstrate that inducing a local immunoinhibitory environment through a single administration of a co-delivered PD-L1 vector is sufficient for durable bNAb expression. To our knowledge, this is the first study to demonstrate a viable strategy that consistently supports sustained concentrations >50 µg mL^−1^ for at least one year. The improved muscle health at the site of administration alone demonstrates the utility of the PD-L1 approach. Future work will need to test the breadth of this approach and continue to evaluate the safety of AAV-expressed immune checkpoint ligands. Nonetheless, our results establish a foundation for durable, protective antibody expression in NHPs and point to a new translational direction for antibody-based HIV prevention and therapy.

## Materials and Methods

### Rhesus macaques

Eighteen Indian-origin rhesus macaques (11 males, 7 females), aged 3.2–5.0 years at the time of AAV administration, were enrolled in this study. Macaques were housed at the Emory National Primate Research Center (ENPRC) in Atlanta, Georgia. All protocols were approved by the Emory University Institutional Animal Care and Use Committee (Permit number: 202100453) and were performed in accordance with guidelines established by the United States Department of Agriculture (USDA) Animal Welfare Act and the recommendations of the National Institutes of Health (NIH) Guide for the Care and Use of Laboratory Animals (8th Edition). The animal care facilities at ENPRC are accredited by both the USDA and AAALAC, International. Macaques were housed in pairs with compatible animals for the entirety of the study. Macaques were screened for serum neutralizing antibodies against the AAV9 serotype prior to study initiation. Animals were stratified into five groups based on age, weight, sex, and baseline serum AAV9 neutralizing antibody (NAb) titers. Three macaques in the PD-L1-only group (Mm013, Mm014, Mm017) had serum AAV9 ID_50_ titers between 1:5 to 1:20; all other macaques had ID_50_ titers <1:5. Body weights at the time of vector administration ranged from 5.3–9.3 kg. All macaques were SIV-negative and *Mamu-B*08*⁻/*Mamu-B*17*⁻. Macaque identifiers, *Mamu-A*01* and *Mamu-A*02* genotype, sex, weight, and age at study initiation are detailed in **Table S1**.

### Cell Lines and Plasmids

HEK293T cells were purchased from ATCC and cultured in DMEM supplemented with 10% fetal bovine serum (FBS), 0.5% penicillin-streptomycin, and 10 mM HEPES, and maintained at 37°C with 5% CO₂. Expi293F cells (Gibco) were cultured in Expi293 Expression Medium (Gibco) according to the manufacturer’s instructions. Expression plasmids used for pseudovirus production for pNL4-3Δenv, BG505, and SIVmac239 have been previously described (*29, 30*). The following reagents were obtained from the NIH AIDS Reagent Program: TZM-bl cells (HRP-8129, contributed by Dr. John C. Kappes, Dr. Xiaoyun Wu, and Tanzyme, Inc.) TRO11, CNE8, BJOX2000, X1632, CE1176, CH119, CE0217, CNE55 (cat# 12670, contributed by Dr. David Montifiori); PVO.4 (ARP-11022, contributed by Dr. David Montefiori, Dr. Feng Gao, and Dr. Ming Li); 9014 (ARP-11571, contributed by Dr. Beatrice H. Hahn, Dr. Brandon F. Keele, and Dr. George M. Shaw); RSC3 expression plasmid (ARP-12279, contributed by Dr. Zhi-Yong Yang, Dr. Gary Nabel, and Dr. John Mascola). The transfer plasmid for pAAV.CAG.fLuc was a gift from Dr. Mark Kay (Addgene # 83281). The expression plasmid encoding furin was previously described (*10*). The SHIV_AD8-EO_ molecular clone was a gift from Dr. Malcolm Martin.

### AAV Production and Purification

AAV 3BNC117 and PD-L1 transgenes were synthesized by GenScript. All transgenes were codon-optimized and cloned into an AAV transfer plasmid including the AAV serotype 2 inverted terminal repeats (AAV2-ITRs) using NotI restriction sites. Production of recombinant AAV was performed by the University of Massachusetts Medical School Vector Core and has been previously described (*31*). Briefly, HEK293T cells were transfected with 1 of 3 AAV transfer plasmids used in this study, a plasmid encoding AAV2 rep and the AAV9 capsid, and a helper plasmid encoding adenovirus genes. Following collection of transfected cell lysates, AAV9 vectors were purified using three consecutive CsCl centrifugation steps. The vector genome (vg) copy number was determined by qPCR. AAV particle quality was assessed by electron microscopy, and preparation purity was confirmed by silver-stained SDS-PAGE.

### AAV vector administration

Macaques were administered AAV9.PD-L1 and/or AAV9.3BNC117 at a dose of 2.5 × 10¹² vg kg⁻¹ of each vector in PBS at a volume less than 1 mL. The vectors were delivered intramuscularly across eight sites: two injections in each of the lower and upper quadriceps, biceps, and deltoid muscles.

### Protein Production and Purification

For production of recombinant 3BNC117, HEK293T cells in T225 flasks were transfected with a 3BNC117-encoding AAV transfer plasmid and furin-encoding plasmid using PEIpro transfection reagent (Polyplus) according to the manufacturer’s instructions. DMEM was replaced after 24 h with FreeStyle 293 Expression Medium (Invitrogen). Media was collected 48 h later, clarified by centrifugation at 4,500 × g for 10 min and filtered through 0.45 µm filter flasks (Thermo Scientific). Antibody was purified using HiTrap MabSelect SuRe columns (Cytiva) and eluted with IgG Elution Buffer (Thermo Scientific) into 1 M Tris-HCl (pH 8). Buffer exchange into PBS was performed using Amicon Ultra centrifugal filter tubes (Sigma). Heavy and light chain composition was assessed by Coomassie-stained SDS-PAGE. For RSC3 production, Expi293F cells (Gibco) were diluted to 3 × 10⁶ cells ml⁻¹ and transfected with FectoPRO reagent (Polyplus) according to the manufacturer’s instructions. After 5 days, supernatants were clarified by centrifugation at 4,500 × g for 10 min and filtered through 0.45 µm filter flasks. Imidazole was added to clarified supernatants to a final concentration of 10 mM. Proteins were purified using HisTrap HP columns (Cytiva) and eluted with 300 mM imidazole. Eluates were concentrated and buffer-exchanged into PBS using Amicon Ultra centrifugal filter tubes (Sigma). Recombinant proteins were aliquoted and stored at 4°C for short-term use.

### Challenge virus production and challenge procedure

Preparation of the R5-tropic, tier-2 SHIV_AD8-EO_ stock derived from rhesus macaque PBMCs has been previously described (*32*). The 10 TCID₅₀ challenge dose, equivalent to 0.27 animal infectious doses (AID₅₀), was selected based on prior studies (*15*). Challenge virus was diluted in 1.0 mL of serum-free RPMI medium to 10 TCID_50_, after which it was loaded into a 3 mL syringe and delivered intrarectally to each macaque. Beginning at 20-weeks post AAV administration, macaques were challenged every two weeks for up to 10 total exposures or until infection was confirmed twice by qRT-PCR.

### SHIVgag Plasma Viral Load Quantification

Quantification of the plasma viral load was performed as previously described (*33*). Briefly, the QIAsymphony DSP Virus/Pathogen Mini Kit (Qiagen) was used to extract viral RNA from rhesus macaque plasma, according to the manufacturer’s recommendations, with a 200 µL input volume and 60 µL elution volume. Quantification of SHIV RNA was performed by one-step qRT-PCR using the TaqMan Fast Virus 1-Step Master Mix (Applied Biosystems) and SHIV/SIVgag-specific primers and probe (forward: 5′-GCAGAGGAGGAAATTACCCAGTAC-3′, Fisher Scientific; reverse: 5′-CAATTTTACCCAGGCATTTAATGTT-3′, Fisher Scientific; probe: 5′-6FAM-TGTCCACCTGCCATTAAGCCCGA-TAMRA-3′, Applied Biosystems). A standard curve was generated from five-fold serial dilutions of SHIV/SIVgag plasmid RNA ranging from 2.11 × 10^8^ to 2.69 × 10^3^ copies mL^−1^, which were run in duplicate on each plate. Reactions were performed on the Applied Biosystems 7500 or QuantStudio 3 Real-Time PCR System using the following cycling profile: 50°C for 15 min, 95°C for 2 min, and 40 cycles of 95°C for 15 s and 60°C for 1 min. Viral RNA concentrations (copies mL^−1^) of plasma were calculated from the standard curve (R² ≥ 0.99) and adjusted for sample dilution if applicable. The assay limit of detection is 60 copies mL^−1^.

### TZM-bl Neutralization assay

HIV-1, SHIV, and SIV pseudoviruses were produced as previously described (*29, 30, 34–36*). Heat inactivated sera were diluted 1:5 in DMEM and then serially diluted five-fold, yielding final assay dilutions from 1:10 to 1:31,250. Each dilution was tested in duplicate in a 96-well plate. Diluted sera were then mixed 1:1 (v/v) with pseudoviruses in DMEM and incubated at 37°C for 30 min. Next, 10,000 TZM-bl cells were added to each well. After incubation at 37°C for 48 h, luciferase activity was determined using Britelite Plus (Revvity) and read on a BioTek Synergy Neo2 plate reader (Agilent). ID₅₀ values were determined by fitting the data to a four-parameter logistic regression model.

### gp120 and ADA ELISAs

Costar 96-well half-area assay plates were coated overnight at 4°C with 3 µg mL^−1^ gp120-CNE8 (Immune Technology) in PBS to determine serum concentrations of 3BNC117. Additional ELISAs were performed using 6 µg mL^−1^ purified RSC3 to determine post-challenge 3BNC117 concentrations and 3 µg mL^−1^ of 3BNC117 antibody to assess ADA responses. A blocking buffer consisting of 5% skim milk, 5% bovine serum albumin (BSA) (Fisher Scientific), and 0.1% Tween-20 was used for blocking the ELISA plates, sample dilutions, and secondary antibody dilutions. Plates were washed twice with PBS-T (PBS with 0.05% Tween-20) and then blocked with blocking buffer for 1 h at 37°C. Sera were heat inactivated at 56°C for 30 min, followed by the addition of 0.1% Tween-20. Sera were then serially diluted in the blocking buffer and added to the plate in duplicate. Purified 3BNC117 was used to generate standard curves as appropriate. Samples were incubated at 37°C for 1 h, then plates were washed five times with PBS-T. For assessing bNAb concentration, a 1:5000 dilution of a horseradish peroxidase (HRP)-conjugated antibody targeting the IgG Fc (Jackson Immuno Research) was added. For ADA ELISAs, a 1:8000 dilution of an HRP-conjugated anti-human lambda light chain antibody (Millipore Sigma) was used to quantify 3BNC117 binding antibodies. The plates were then incubated at 37°C for 1 h, after which they were washed ten times with PBS-T. 3,3’,5,5’-Tetramethylbenzidine (TMB) Substrate Solution (Thermo Fisher Scientific) was added to the plates. The reaction was stopped by the addition of TMB Stop Solution (KPL) between 2-10 min depending on the assay. Absorbance was measured at 450 nm using a BioTek Synergy Neo2 plate reader (Agilent). Endpoint titers determined by identifying the serum dilution that produced an optical density (O.D.) of 0.2.

### AAV neutralization assay

AAV neutralization assays were performed as previously described (*37, 38*). Heat inactivated sera were diluted 1:5 in DMEM and then serially diluted four-fold, yielding final assay dilutions from 1:10 to 1:5,120. Each dilution was tested in duplicate in a 96-well plate. Diluted sera were then mixed 1:1 (v/v) with 10^10^ vg mL^−1^ of AAV9.CAG.fLuc vector and incubated for 30 min at 37°C. Next, 25,000 HEK293T cells were added to each well. After incubation at 37°C for 24 h, luciferase activity was determined using Britelite Plus (Revvity) and read on a BioTek Synergy Neo2 plate reader (Agilent).

### ELISpot assays

Milipore MultiScreen-HA Filter plates (MAHA S4519; Milipore, MA) were coated with 5 µg/mL mouse anti-human IFN-γ (Pharmingen) in PBS at 4°C overnight then washed four times with RPMI containing 10% FBS and 1% penicillin/streptomycin (FRPMI), and blocked with FRPMI for 1 h. PBMCs previously isolated by density gradient centrifugation using Ficoll-Pacque PLUS (Cytiva) were thawed, washed in FRPMI with 50 U/mL benzonase, resuspended in FRPMI, and rested for at least 2 h at 37°C. Peptide pools were synthesized by GenScript and covered the variable regions of the bNAb heavy (3BNC117 VarH) and light (3BNC117 VarL) chains; the LS-Furin-P2A region; the IgG1 constant heavy chain region; and the kappa light chain constant regions. Appropriately diluted peptide pools, positive-control staphylococcal enterotoxin B (SEB; ToxTech), or DMSO (Sigma) were dispensed into the 96-well plate in duplicate; 200,000–250,000 PBMCs were then added per well to yield final concentrations of 10 µg mL⁻¹ for each peptide pool, 4 µg mL⁻¹ for SEB, and 0.1% DMSO for vehicle controls. Plates were incubated for 20 h, washed, and incubated with 1 µg mL^−1^ biotin-conjugated anti-human IFN-γ mAb in PBS-T + 1% FBS (PBS-T-FBS) for 3 h, followed by a 1:1000 dilution of streptavidin-conjugated alkaline phosphatase (Rockland) in PBS-T-FBS for 1 h at 37 °C. After washing the plate four times with PBS-T, spots were developed with One-Step NBT/BCIP (Thermo Scientific) and quantified using CTL ImmunoSpot 7.0 software (Cellular Technology Ltd).

### H&E and Immunohistochemistry

Muscle samples from rhesus macaques were fixed in 10% neutral buffered formalin (NBF), processed, and blocked in paraffin for histological analysis. All samples were sectioned at 5μm and stained with hematoxylin and eosin (H&E) for routine histopathology. Staining for multiple antibodies, including CD3 (1:200, Abcam, ab16669); CD4 (1:200, Abcam, ab13316); CD19 (1:200, Abcam, ab134114); CD68 (1:200, Thermo Fisher, MA5-13324); FoxP3 (1:200, Abcam, ab20034); CD56 (Leica, CD56-504) and PD-1 (1:200, Sino Biologicals, 90305-MM09) were achieved on the Bond RX automated system with the Bond Polymer Refine Red Detection (DS9390) (Leica) used per manufacturer’s protocol. Tissue sections were dewaxed with Bond Dewaxing Solution (Leica) at 72°C for 30 min. Heat-induced epitope retrieval was performed using Epitope Retrieval Solution (Leica), heated to 100°C for 20 min. Sections were examined under light microscopy using an Olympus BX51 microscope. Photographs were taken using an Olympus DP73 camera and were evaluated by a board-certified veterinary pathologist in a blinded manner.

### Spatial Transcriptomics

Tissue slides were generated, stained and imaged according to instructions provided by 10x Genomics (Tissue Preparation Handbook, CG000684) at the ENPRC Histology and Molecular Pathology Laboratories. Visium HD Spatial Gene Expression (Probe-based) Libraries were constructed using the Visium HD Spatial Gene Expression Reagent Kit (10x Genomics) according to the manufacturer’s instructions at the ENPRC Genomics Core. First, RNA was extracted from scrolls collected from the formalin-fixed, paraffin-embedded (FFPE) tissue blocks with the RNeasy FFPE kit (Qiagen), and subjected to RNA quality assessment, where samples were confirmed to have DV200 scores >30%. Tissue sections with 5 μm thickness were prepared from the FFPE blocks and placed onto glass slides, subjected to H&E staining, and imaged at 40x magnification using a NanoZoomer S60 Digital Slide Scanner C13210 (Hamamatsu). The 6.5 mm x 6.5 mm area of interest (AOI) for each tissue section was then chosen, followed by decrosslinking and hybridization of the Visium Human Transcriptome Probes Set v2.0 with the tissue in these AOIs. The hybridized probes are released and captured by the Visium HD slide oligos using the Visium CytAssist instrument (10x Genomics), amplified, and the final sequencing libraries were generated. Libraries were validated by capillary electrophoresis on a TapeStation 4200 (Agilent) and sequenced with PE100 reads and targeting a depth of 300 million reads per sample on a NovaSeq 6000 (Illumina). The resultant bcl files from sequencing on the NovaSeq 6000 was demultiplexed via bcl2fastq to generate FASTQ files. The Space Ranger v4.0.1 pipeline was used to map FASTQ files to the human GRCh38-2020-A reference and Visium Human Transcriptome Probe Set v2.0, detect the tissue section, align the sequencing data to the microscope image and the CytAssist image, and output gene-barcode matrices for further analysis (*39*). Downstream analysis was performed on the data output from Space Ranger at the 8 μm binned resolution. Individual transcript localization, graph-based clustering figures, and clustered gene expression heat maps were generated in Loupe Browser v9.0.0.

### Statistical analysis

Data were analyzed using GraphPad Prism v10.4 (GraphPad, La Jolla, CA). Comparisons of groups were performed as indicated in manuscript and/or reported in the figures legends with statistical significance reported as a p value ≤ 0.05.

## Acknowledgements

The authors would like to thank M. Farzan, C.C. Bailey, M.A. Martins, and M.D. Alpert for their discussions and insights regarding this study; the Emory National Primate Research Center (ENPRC) staff for their tremendous help in completing this study; and M.E. Davis-Gardner for her comments and edits to the manuscript. This work was supported in part by National Institutes of Health awards R01AI167724 (M.R.G.) and R01DA056770 (M.R.G.). Additional support was provided from the NIH Office of Research Infrastructure Programs (ORIP) P51OD11132 to ENPRC, U42OD011023 to ENPRC, and P30AI050409 to the Emory University Center for AIDS Research. Next generation sequencing services were provided by the Emory NPRC Genomics Core (RRID:SCR_026418) which is supported in part by NIH P51OD011132. Sequencing data was acquired on an Illumina NovaSeq 6000 funded by NIH S10OD026799.

## Author Contributions

MRG conceived the study and acquired funding. MK, AAK, IL, PD, YB, MY-HL, SL, MC, DM, EDD, and MRG performed experiments. SW, CW, SE, RLS, JSW, and EHC were responsible for performing the required procedures and sample collection of the nonhuman primate study. YN provided critical reagents and input into the study design. JX and GG were responsible for production and quality control of the AAV vectors. MK, MYL, GKT, SL, MC, SV, DAK, INM, SEB, and MRG were responsible for data analysis. MK, SEB, and MRG wrote the initial draft of the manuscript which was reviewed and approved by all coauthors.

## Competing Interests

MK, AAK, IL, and MRG are named inventors on a patent application related to the technologies described in this study submitted by Emory University. MRG is a co-founder and consultant for Emmune, Inc. MRG has consulted for ViiV Healthcare. GG is a co-founder of Voyager Therapeutics and Aspa Therapeutics and holds equity in both companies. GG is an inventor on patents with potential royalties licensed to Voyager Therapeutics, Aspa Therapeutics, and other biopharmaceutical companies. The remaining authors declare no competing interests regarding this study.

## Data, code and material availability

All spatial transcriptomic data can be accessed on NCBI GEO, Accession Number: GSE314035.

## Supplementary Materials

### Supplemental Figure Legends

**Fig. S1.**
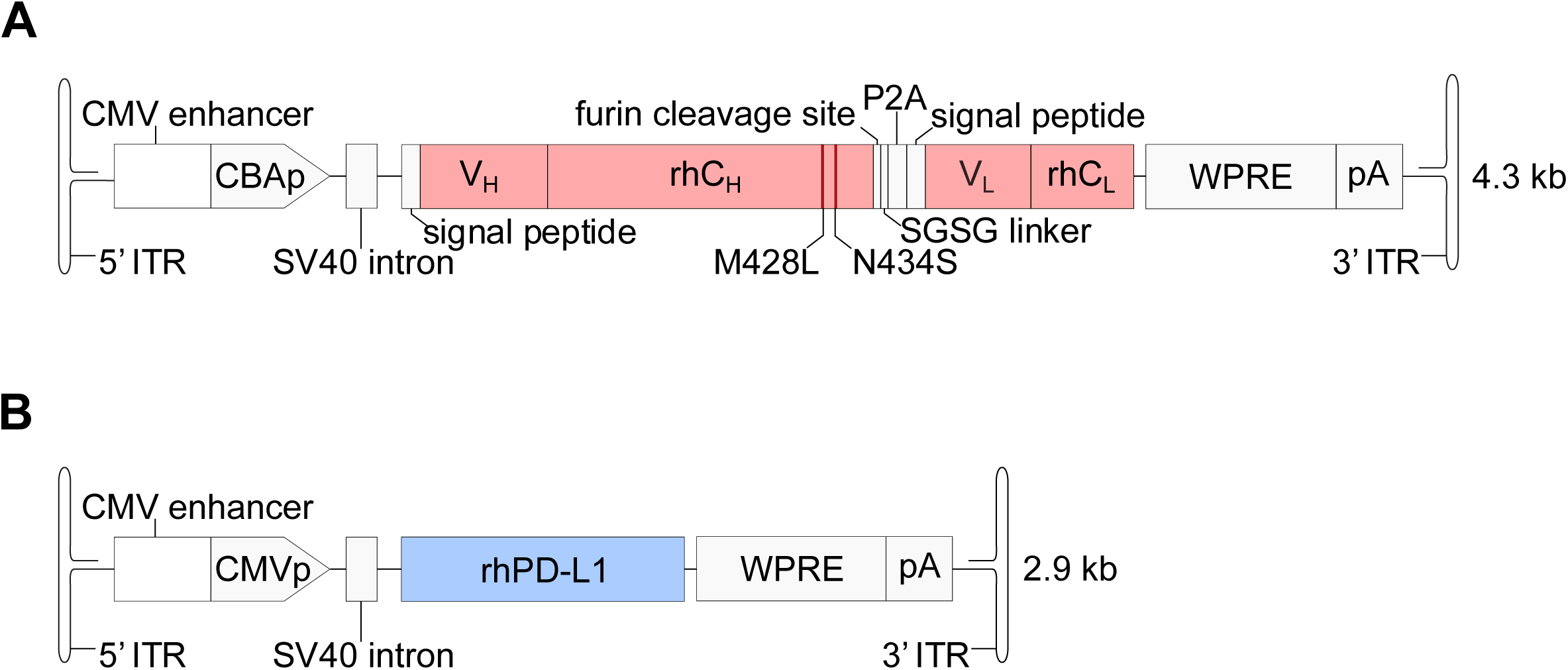
AAV9 vectors encoding 3BNC117 or rhesus PD-L1. **(A)** Diagram of the AAV transgene cassette encoding 3BNC117. 3BNC117 was “rhesusized” by exchanging its native human constant regions for rhesus macaque constant regions. **(B)** Diagram of the AAV transgene cassette encoding PD-L1 (*Macaca mulatta*). Abbreviations: ITR, AAV2 inverted terminal repeat; CMVp, cytomegalovirus immediate-early promoter; CBAp, chicken-beta actin promoter; SV40 intron, simian virus 40 intron; V_H_, variable heavy chain region; rhC_H_. rhesus macaque (rh) constant heavy chain (IgG1); furin cleavage site, RKRR; SGSG linker, serine/glycine linker; P2A, ribosomal skipping peptide from porcine teschovirus-1; V_L_, variable light chain region; rhC_L_, rhesus macaque (rh) kappa light chain constant region; WPRE, woodchuck hepatitis virus posttranscriptional regulatory element; pA, SV40 polyadenylation signal sequence; kb, kilobases. Note the M428L/N434S half-life-extending amino acid substitutions in the rhC_H_ region. The size of the AAV transgene cassettes is indicated on the right.

**Fig. S2.**
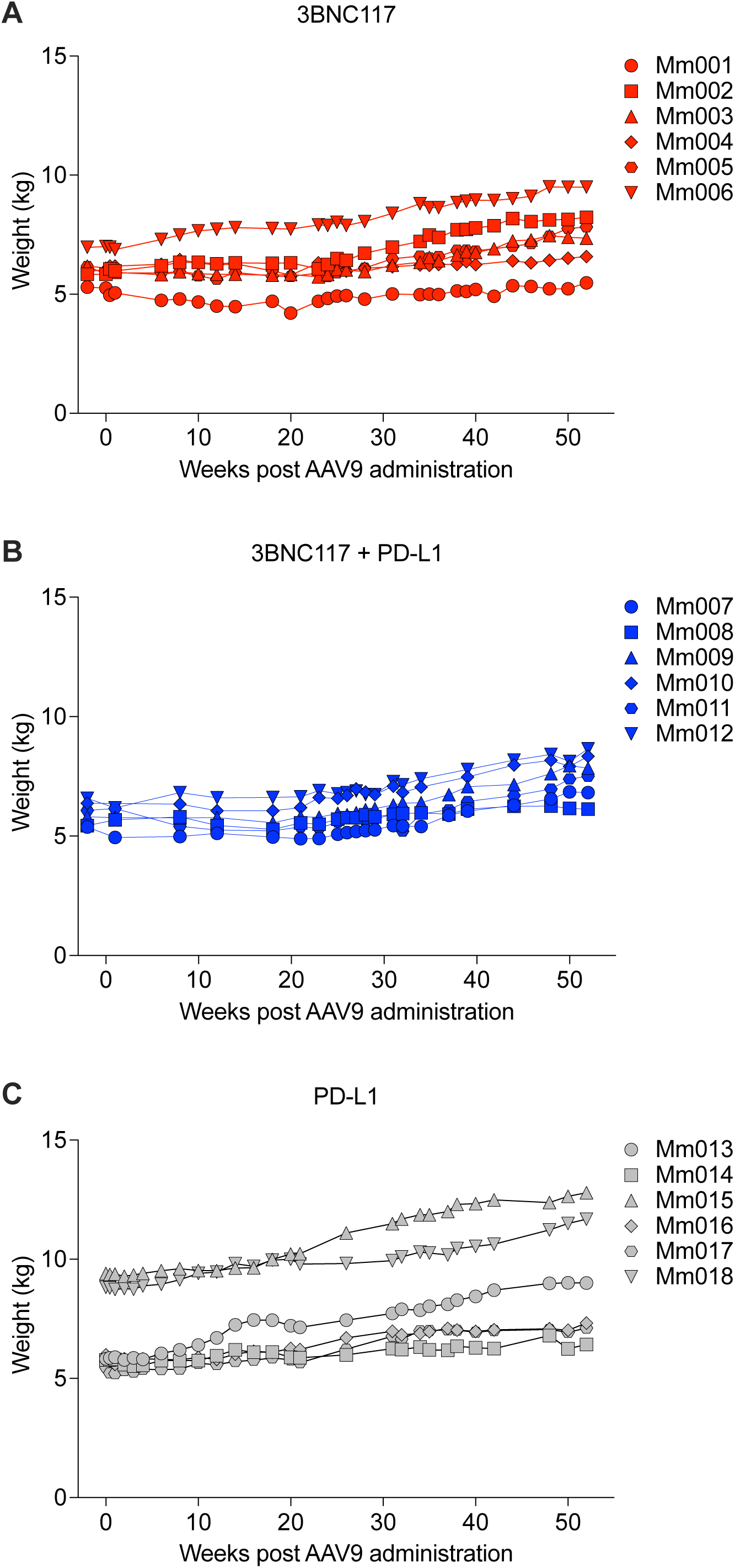
Macaque weight gain over the course of the study. Weight gain over the course of 52 weeks in macaques that received **(A)** AAV9.3BNC117 only, **(B)** AAV9.3BNC117 plus AAV9.PD-L1 or **(C)** AAV9.PD-L1 only.

**Fig. S3.**
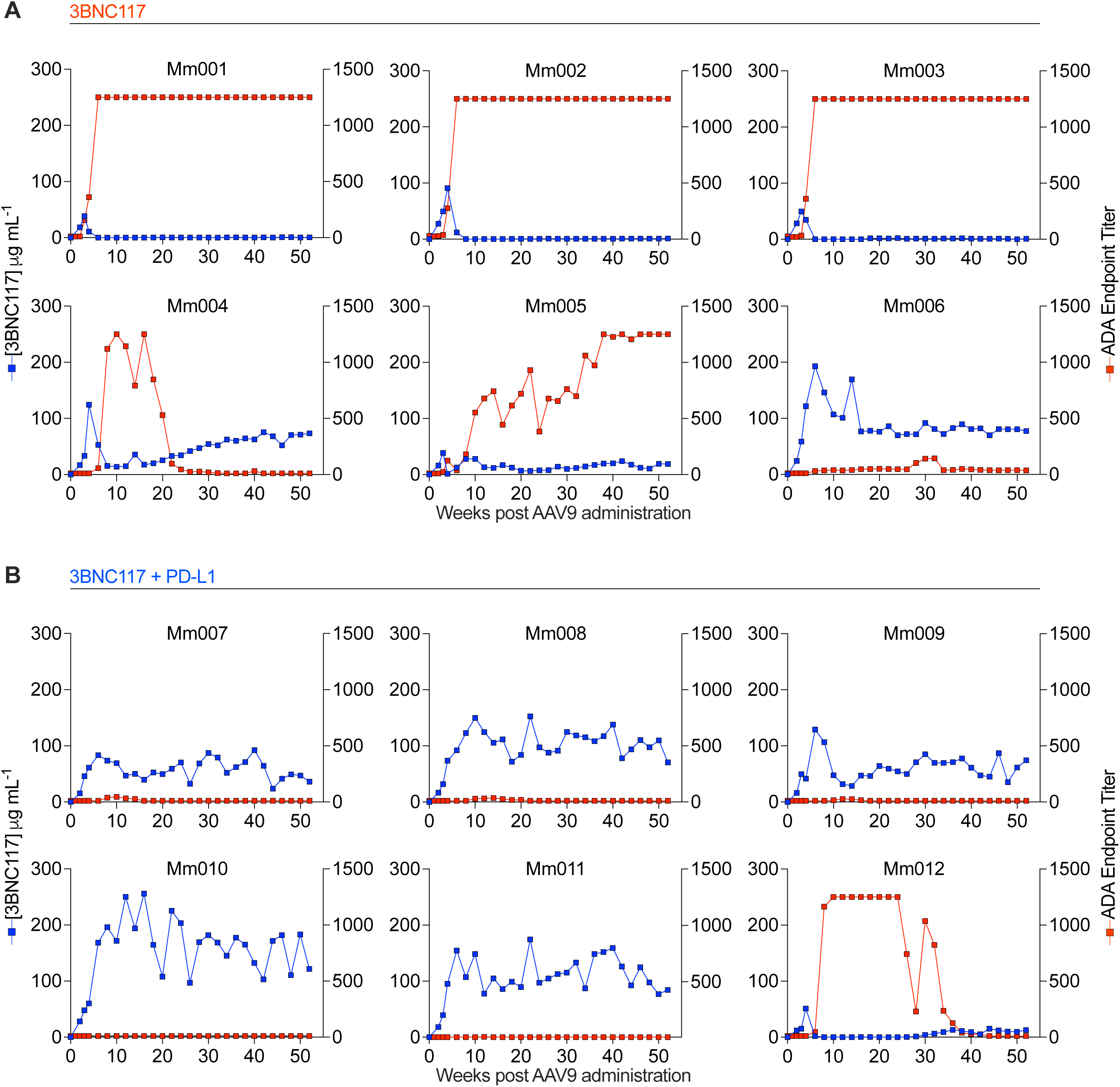
Serum bNAb concentration vs ADA response. Serum bNAb concentrations vs ADA endpoint titers in individual macaques that received **(A)** AAV9.3BNC117 only or **(B)** AAV9.3BNC117 plus AAV9.PD-L1, measured by gp120 ELISA or RSC3 over 52 weeks. For ADA endpoint titers, serum collected 2 weeks prior to study initiation was used as baseline for each animal. ADA endpoint titers are defined as the highest serum dilution with OD₄₅₀ ≥ 0.2.

**Fig. S4.**
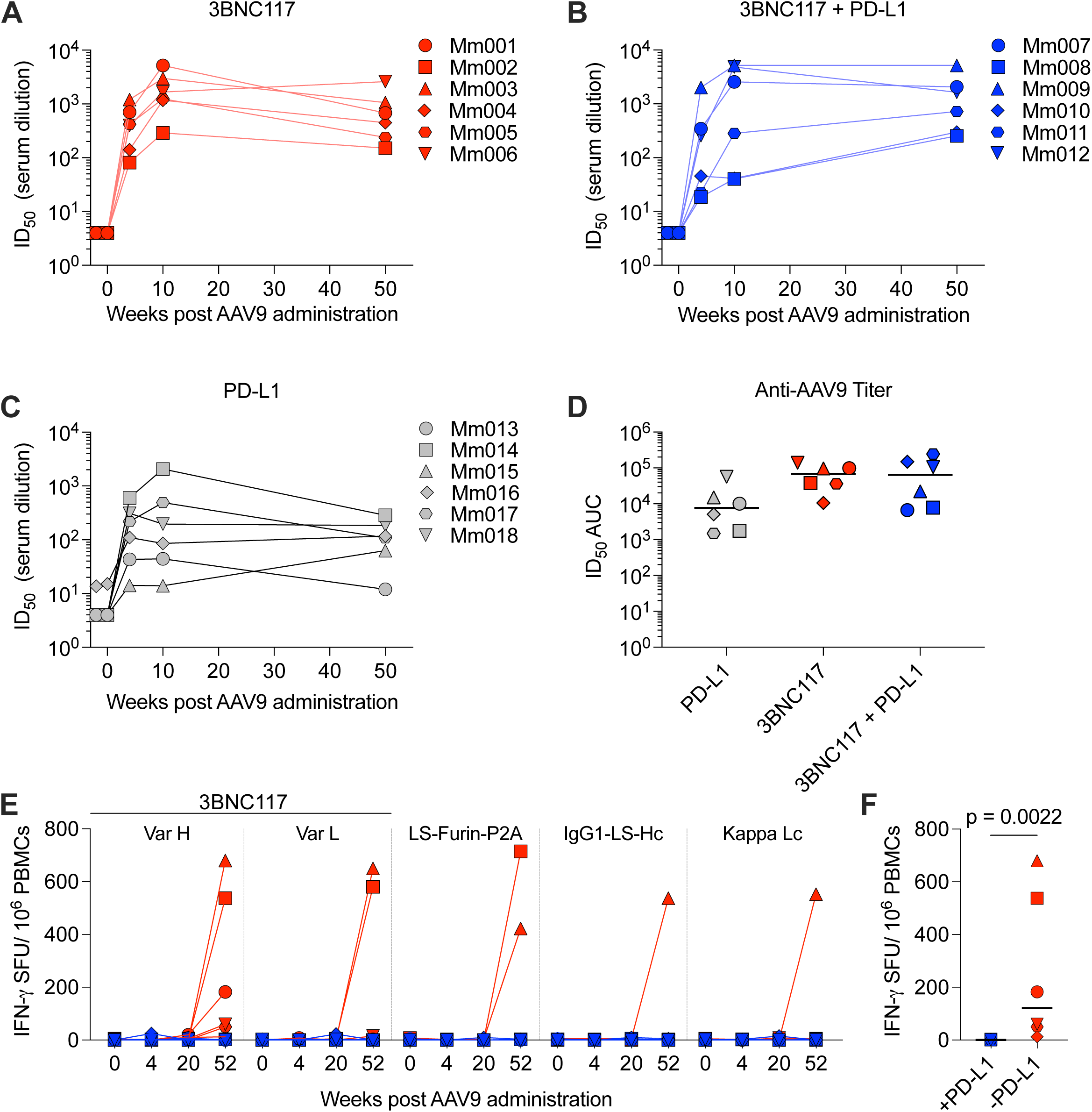
AAV9 neutralizing antibody responses and PBMC IFNγ ELISpot reactivity in macaques. AAV9 neutralization titers of all 18 macaques from Weeks −2, 0, 4, 10 and 50 post AAV9 administration. ID₅₀ values for each macaque per group are reported. Those samples that did not reach 50% neutralization were normalized to a value of <10. Serum collected 2 weeks prior to study initiation was used as baseline for each macaque. Serum ID₅₀ titers in individual macaques that received **(A)** AAV9.3BNC117 only, **(B)** AAV9.3BNC117 plus AAV9.PD-L1 or **(C)** AAV9.PD-L1 only. **(D)** ID_50_ AUC values for data in (A–C). Black lines indicate the median. Note symbols in (D) match those used in panels (A–C), respectively. **(E)** IFNγ ELISpot reactivity for Weeks 0, 4, 20 and 52 PBMC samples against AAV9.3BNC117 peptide pools (3BNC117 VarH, 3BNC117 VarL, LS-Furin-P2A peptide, IgG1 constant heavy chain region, and kappa light chain constant regions) in macaques that received AAV9.3BNC117. **(F)** Comparison of PBMC IFNγ ELISpot reactivity against AAV9.3BNC117 peptide pools at week 52 in macaques that received AAV9.3BNC117 only or AAV9.3BNC117 and AAV9.PD-L1. Black lines indicate the median. Note symbols in (E) and (F) match those used in panels (A) and (B), respectively. Statistical significance in (F) was determined by two-tailed Mann-Whitney test and defined as p ≤ 0.05.

**Fig. S5.**
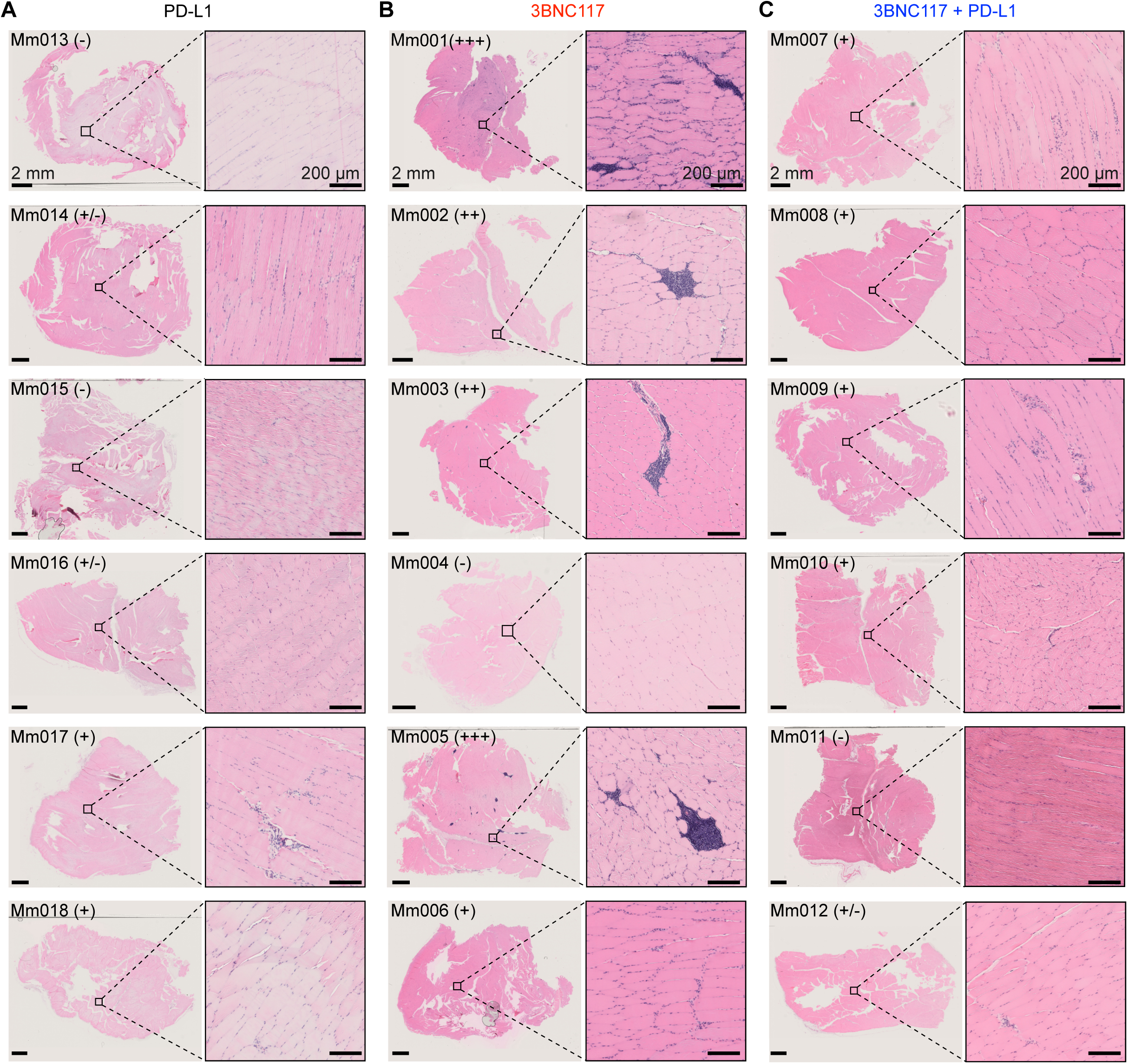
Histopathology staining for macaques that received AAV9 administered vectors. Representative H&E-stained muscle tissue from the upper left quadriceps of all 18 macaques that was harvested at necropsy. Stained tissue from individual macaques that received **(A)** AAV9.PD-L1 only, **(B)** AAV9.3BNC117 only, or **(C)** AAV9.3BNC117 plus AAV9.PD-L1. Regions of interest are enlarged on the right. Scale bars represent 1 mm on the left and 100 µm on the right.

**Fig. S6.**
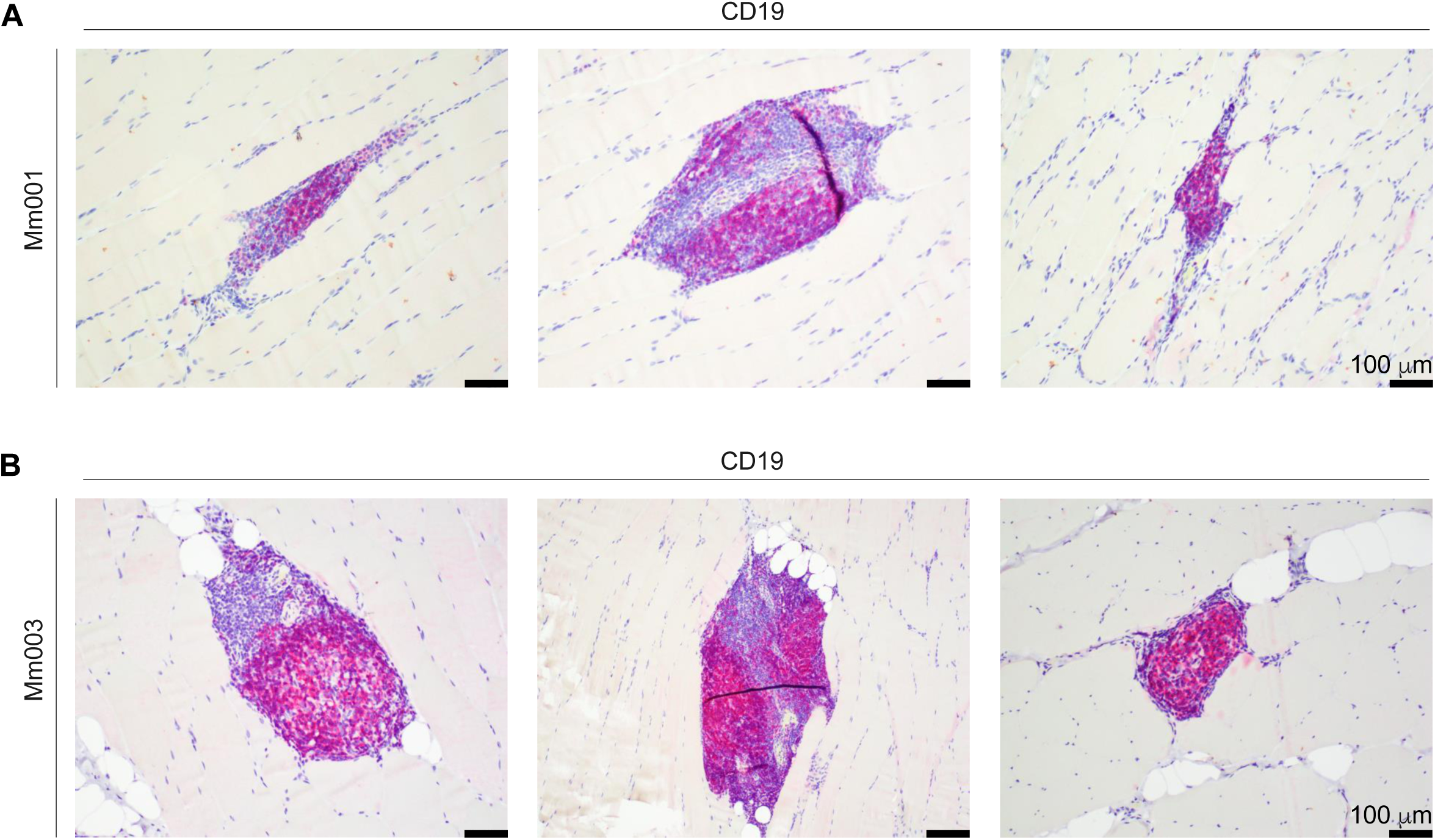
TLSs develop in macaques with severe inflammation at the site of AAV administration. IHC staining for CD19 in **(A)** Mm001 and **(B)** Mm003 showing TLSs. Scale bars represent 100 µm.

**Table S1.**
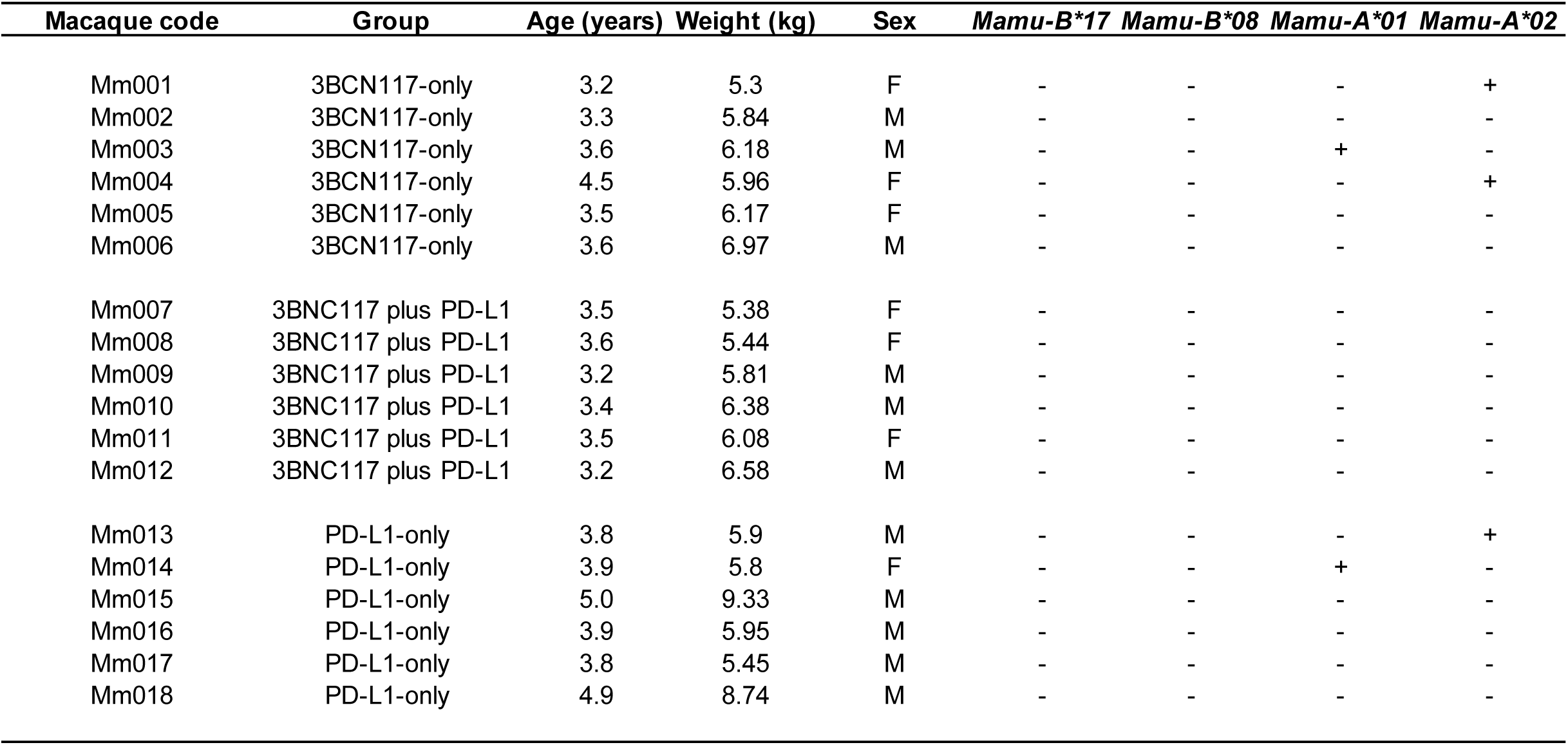
Characteristics of the 18 rhesus macaques enrolled in this study.

| **Macaque code** | **Group** | **Age (years)** | **Weight (kg)** | **Sex** | ***Mamu-B\*17*** | ***Mamu-B\*08*** | ***Mamu-A\*01*** | ***Mamu-A\*02*** |
| --- | --- | --- | --- | --- | --- | --- | --- | --- |
| Mm001 | 3BCN117-only | 3.2 | 5.3 | F | - | - | - | + |
| Mm002 | 3BCN117-only | 3.3 | 5.84 | M | - | - | - | - |
| Mm003 | 3BCN117-only | 3.6 | 6.18 | M | - | - | + | - |
| Mm004 | 3BCN117-only | 4.5 | 5.96 | F | - | - | - | + |
| Mm005 | 3BCN117-only | 3.5 | 6.17 | F | - | - | - | - |
| Mm006 | 3BCN117-only | 3.6 | 6.97 | M | - | - | - | - |
| Mm007 | 3BNC117 plus PD-L1 | 3.5 | 5.38 | F | - | - | - | - |
| Mm008 | 3BNC117 plus PD-L1 | 3.6 | 5.44 | F | - | - | - | - |
| Mm009 | 3BNC117 plus PD-L1 | 3.2 | 5.81 | M | - | - | - | - |
| Mm010 | 3BNC117 plus PD-L1 | 3.4 | 6.38 | M | - | - | - | - |
| Mm011 | 3BNC117 plus PD-L1 | 3.5 | 6.08 | F | - | - | - | - |
| Mm012 | 3BNC117 plus PD-L1 | 3.2 | 6.58 | M | - | - | - | - |
| Mm013 | PD-L1-only | 3.8 | 5.9 | M | - | - | - | + |
| Mm014 | PD-L1-only | 3.9 | 5.8 | F | - | - | + | - |
| Mm015 | PD-L1-only | 5.0 | 9.33 | M | - | - | - | - |
| Mm016 | PD-L1-only | 3.9 | 5.95 | M | - | - | - | - |
| Mm017 | PD-L1-only | 3.8 | 5.45 | M | - | - | - | - |
| Mm018 | PD-L1-only | 4.9 | 8.74 | M | - | - | - | - |

## References and Notes

1. Y. Bar-On et al., Safety and antiviral activity of combination HIV-1 broadly neutralizing antibodies in viremic individuals. Nat Med 24, 1701–1707 (2018).

2. P. Mendoza et al., Combination therapy with anti-HIV-1 antibodies maintains viral suppression. Nature 561, 479–484 (2018).

3. L. Corey et al., Two Randomized Trials of Neutralizing Antibodies to Prevent HIV-1 Acquisition. N Engl J Med 384, 1003–1014 (2021).

4. C. Gaebler et al., Prolonged viral suppression with anti-HIV-1 antibody therapy. Nature 606, 368–374 (2022).

5. M. C. Sneller et al., Combination anti-HIV antibodies provide sustained virological suppression. Nature 606, 375–381 (2022).

6. J. M. Martinez-Navio et al., Long-Term Delivery of an Anti-SIV Monoclonal Antibody With AAV. Front Immunol 11, 449 (2020).

7. J. M. Martinez-Navio et al., Host Anti-antibody Responses Following Adeno-associated Virus-mediated Delivery of Antibodies Against HIV and SIV in Rhesus Monkeys. Mol Ther 24, 76–86 (2016).

8. S. P. Fuchs et al., AAV-Delivered Antibody Mediates Significant Protective Effects against SIVmac239 Challenge in the Absence of Neutralizing Activity. PLoS Pathog 11, e1005090 (2015).

9. K. O. Saunders et al., Broadly Neutralizing Human Immunodeficiency Virus Type 1 Antibody Gene Transfer Protects Nonhuman Primates from Mucosal Simian-Human Immunodeficiency Virus Infection. J Virol 89, 8334–8345 (2015).

10. M. R. Gardner et al., Anti-drug Antibody Responses Impair Prophylaxis Mediated by AAV-Delivered HIV-1 Broadly Neutralizing Antibodies. Mol Ther 27, 650–660 (2019).

11. J. M. Martinez-Navio et al., Adeno-Associated Virus Delivery of Anti-HIV Monoclonal Antibodies Can Drive Long-Term Virologic Suppression. Immunity 50, 567–575.e565 (2019).

12. F. H. Priddy et al., Adeno-associated virus vectored immunoprophylaxis to prevent HIV in healthy adults: a phase 1 randomised controlled trial. Lancet HIV 6, e230–e239 (2019).

13. J. P. Casazza et al., Safety and tolerability of AAV8 delivery of a broadly neutralizing antibody in adults living with HIV: a phase 1, dose-escalation trial. Nat Med 28, 1022–1030 (2022).

14. G. J. Freeman et al., Engagement of the PD-1 immunoinhibitory receptor by a novel B7 family member leads to negative regulation of lymphocyte activation. J Exp Med 192, 1027–1034 (2000).

15. R. Gautam et al., A single injection of crystallizable fragment domain-modified antibodies elicits durable protection from SHIV infection. Nat Med 24, 610–616 (2018).

16. S. Khanal, A. Wieland, A. J. Gunderson, Mechanisms of tertiary lymphoid structure formation: cooperation between inflammation and antigenicity. Front Immunol 14, 1267654 (2023).

17. A. Kratz, A. Campos-Neto, M. S. Hanson, N. H. Ruddle, Chronic inflammation caused by lymphotoxin is lymphoid neogenesis. J Exp Med 183, 1461–1472 (1996).

18. M. Bombardieri et al., Inducible tertiary lymphoid structures, autoimmunity, and exocrine dysfunction in a novel model of salivary gland inflammation in C57BL/6 mice. J Immunol 189, 3767–3776 (2012).

19. L. G. M. Heezen et al., Spatial transcriptomics reveal markers of histopathological changes in Duchenne muscular dystrophy mouse models. Nat Commun 14, 4909 (2023).

20. S. Adriouch et al., Improved Immunological Tolerance Following Combination Therapy with CTLA-4/Ig and AAV-Mediated PD-L1/2 Muscle Gene Transfer. Front Microbiol 2, 199 (2011).

21. P. O. Carlsson et al., Survival of Transplanted Allogeneic Beta Cells with No Immunosuppression. N Engl J Med 393, 887–894 (2025).

22. T. B. McMurphy et al., AAV-mediated co-expression of an immunogenic transgene plus PD-L1 enables sustained expression through immunological evasion. Sci Rep 14, 28853 (2024).

23. V. A. Klenchin et al., Adeno-associated viral delivery of Env-specific antibodies prevents SIV rebound after discontinuing antiretroviral therapy. Sci Immunol 10, eadq4973 (2025).

24. S. P. Fuchs et al., Transient rapamycin treatment avoids unwanted host immune responses toward AAV-delivered anti-HIV antibodies. Nat Commun 16, 8906 (2025).

25. J. R. Mendell et al., Sustained alpha-sarcoglycan gene expression after gene transfer in limb-girdle muscular dystrophy, type 2D. Ann Neurol 68, 629–638 (2010).

26. O. Cao et al., Induction and role of regulatory CD4+CD25+ T cells in tolerance to the transgene product following hepatic in vivo gene transfer. Blood 110, 1132–1140 (2007).

27. S. P. Fuchs, J. M. Martinez-Navio, E. G. Rakasz, G. Gao, R. C. Desrosiers, Liver-Directed but Not Muscle-Directed AAV-Antibody Gene Transfer Limits Humoral Immune Responses in Rhesus Monkeys. Mol Ther Methods Clin Dev 16, 94–102 (2020).

28. A. Ardeshir et al., Determinants of successful AAV-vectored delivery of HIV-1 bNAbs in early life. Nature 645, 1020–1028 (2025).

29. M. R. Gardner et al., CD4-Induced Antibodies Promote Association of the HIV-1 Envelope Glycoprotein with CD4-Binding Site Antibodies. J Virol 90, 7822–7832 (2016).

30. C. H. Fellinger, M. R. Gardner, C. C. Bailey, M. Farzan, Simian Immunodeficiency Virus SIVmac239, but Not SIVmac316, Binds and Utilizes Human CD4 More Efficiently than Rhesus CD4. J Virol 91, (2017).

31. C. Mueller, D. Ratner, L. Zhong, M. Esteves-Sena, G. Gao, Production and discovery of novel recombinant adeno-associated viral vectors. Curr Protoc Microbiol Chapter 14, Unit14D.11 (2012).

32. M. Shingai et al., Most rhesus macaques infected with the CCR5-tropic SHIV(AD8) generate cross-reactive antibodies that neutralize multiple HIV-1 strains. Proc Natl Acad Sci U S A 109, 19769–19774 (2012).

33. G. Silvestri et al., Nonpathogenic SIV infection of sooty mangabeys is characterized by limited bystander immunopathology despite chronic high-level viremia. Immunity 18, 441–452 (2003).

34. M. R. Gardner et al., AAV-expressed eCD4-Ig provides durable protection from multiple SHIV challenges. Nature 519, 87–91 (2015).

35. M. R. Gardner et al., AAV-delivered eCD4-Ig protects rhesus macaques from high-dose SIVmac239 challenges. Sci Transl Med 11, (2019).

36. M. E. Davis-Gardner et al., A strategy for high antibody expression with low anti-drug antibodies using AAV9 vectors. Front Immunol 14, 1105617 (2023).

37. M. R. Gardner et al., High concordance of ELISA and neutralization assays allows for the detection of antibodies to individual AAV serotypes. Mol Ther Methods Clin Dev 24, 199–206 (2022).

38. S. Verma et al., Seroprevalence of Adeno-Associated Virus Neutralizing Antibodies in Males with Duchenne Muscular Dystrophy. Hum Gene Ther 34, 430–438 (2023).

39. M. F. Oliveira et al., High-definition spatial transcriptomic profiling of immune cell populations in colorectal cancer. Nat Genet 57, 1512–1523 (2025).

